# Spreading depressions and periinfarct spreading depolarizations in the context of cortical plasticity

**DOI:** 10.1101/2020.01.31.927848

**Authors:** Maria Sadowska, Clemens Mehlhorn, Władysław Średniawa, Łukasz M. Szewczyk, Aleksandra Szlachcic, Paulina Urban, Maciej Winiarski, Jan A. Jabłonka

## Abstract

Studies of cortical function-recovery require a comparison between normal and post-stroke conditions that lead to changes in cortical metaplasticity. Focal cortical stroke impairs experience-dependent plasticity (ExDP) in the neighboring somatosensory cortex and usually evokes periinfarct depolarizations (PiDs) – a spreading depression-like waves. Experimentally induced spreading depressions (SDs) affect gene expression and some of these changes persist for at least 30 days. However, such changes are not stroke-specific: migraine patients had prolonged protein changes after a single SD episode during migraine aura. This study investigates whether non-stroke depolarizations impair cortical ExDP similarly to the stroke.

ExDP was induced in rates with SDs or PiDs by a month of contralateral partial whiskers deprivation. Cortical activity was mapped by [^14^C]-2-deoxy-D-glucose (2DG) incorporation during stimulation of spared and contralateral homotopic whiskers. We found that whiskers deprivation after SDs resulted in normal cortical representation enlargement suggesting that SDs and PiDs depolarization have no influence on ExDP cortical map reorganization. PiDs and the MMP-9, −3, −2 or COX-2 proteins which are assumed to influence metaplasticity in rats after stroke were compared between the SDs induced by high osmolarity KCl solution and the PiDs following cortical photothrombotic stroke (PtS). We found that none of these factors directly caused cortical post-stroke metaplasticity changes. The only significant difference between stoke and induced SD was a greater imbalance in interhemispheric activity equilibrium after stroke. The interhemispheric interactions modified by stroke may therefore be a promising target for future studies of post-stroke ExDP and for convalescence studies.

**Highlights:**

- Post-stroke metaplasticity changes in an injured hemisphere are not a consequence of spreading depolarizations
- None of the monitored proteins (MMP-2, -3, -9; COX-2) cause modifications in poststroke cortical plasticity
- Spreading depressions have a prolonged, global influence on the functioning of both hemispheres and of the thalamus
- Impaired interhemispheric interactions may underlie the post-stroke metaplasticity changes in the injured hemisphere

## Introduction

The primary somatosensory cortex of the whiskers — the barrel field (BF) — is a good model for studying cortical plasticity due to its well-defined somatotopic organization. The whiskers cortical representation is readily rearranged in response to sensory input modifications. In mice, a cortical representation of a row of whiskers, functionally visualized by 2DG incorporation, is enlarged in width by more than 50% when the surrounding whiskers are cut for 7 days (Dietrich et al. 1985; Kaliszewska et al., 2012). In rats, the same area is enlarged by 40% after one month of whisker-cutting (Jabłonka et al., 2007).

Cortical map reorganization involves CaMKII (Calcium/calmodulin-dependent protein kinase II) and requires NMDA (N-methyl-D-aspartate) receptor-dependent induction of long-term potentiation (LTP, Glazewski et al., 2000; Footitt and Newberry, 1998). It is also accompanied by depressions, similar to long-term depressions (LTD), which weaken synaptic connections within the deprived cortex (Drew and Feldman 2009; Li et al., 2009).

*In vivo* studies after SDs showed that potentiation was uncorrelated with meaningful input to the studied cortex area (Guiou et al., 2005; Piilgaard and Lauritzen, 2009; Faraguna et al., 2010; de Souza et al., 2011). Changes typical for cortical plasticity and cortical map remodeling were induced by SDs in the sensory cortex of rats (Theriot et al., 2012). After stroke, synaptic plasticity seemed to be boosted as well (Hagemann et al., 1998).

We previously found that focal stroke strongly moderates the cortical map rearrangement induced by partial whiskers deprivation (Jabłonka et al., 2012). Therefore, assuming that the molecular and electrophysiological markers of synaptic plasticity are enhanced after stroke, while the ExDP manifestation is reduced, we hypothesized that features we should concentrate on are those that might be correlated to rearrangement of dendritic branches and spines (Tailby et al., 2005).

Marik et al (2010) showed that within days after deprivation was initiated, axons on both excitatory and inhibitory neurons in the spared whiskers sensory representation, elongated in the direction of the neighboring cortex deprived of sensory input. MMP-9 was shown to determine spines’ thinning and elongation (Michaluk et al., 2011), supporting the conjecture that it participates in cortical map rearrangement which occurs through ExDP (Kaliszewska et al., 2012). It was also demonstrated that the contralateral hemisphere is involved in spared whiskers representation-widening in 2DG functional brain mapping (Jabłonka, 2012; Jabłonka et al., 2012; Jabłonka et al., 2014).

A considerable body of data demonstrates that a tiny focal brain injury results in alterations within the entire brain (Witte et al., 2000). A focal cortical stroke changes the brain metabolism for at least few months (Jabłonka et al., 2009; Nonose et al., 2018), and the overall activity in the ipsilateral hemisphere decreases for several days. Following diaschisis, an almost immediate post-stroke increase of metabolism related to spontaneous activity is observed in the contralateral hemisphere (Andrews 1991; Mohajerani et al., 2011). GABA-ergic inhibition seems to be reduced bilaterally while the NMDA and AMPA-dependent excitability increases in the lesioned hemisphere (Que et al., 1999; Schiene et al., 1996; Redecker et al., 2002). Consequently, cortical inhibition bilaterally decreases while excitability increases (Neumann-Haefelin and Witte, 2000). Just before necrotic cell death induction the excitability in the nervous tissue is decreased, and this is followed by an enormous excitation due to hypoxia that leads to excitotoxicity and spreading PiDs (Heiss, 2011). The PiDs systematically cross the entire hemisphere, activating the glial cells and modifying tissue excitability and metabolism (Urbach et al., JCBFM 2014). For example, the growth of contralateral branches was observed as a result of local thermal coagulation lesion (Carmichael and Chesselet 2002).

Our previous studies showed that focal cortical photothrombotic stroke (PtS) in a vicinity of the BF, inhibits the cortical map remodeling by ExDP induced by whiskers deprivation (Cybulska-Kłosowicz et al. 2011; Jabłonka et al., 2007). The inhibition depended on MMPs and COX-2 expression, probably reflecting an inflammatory response (Cybulska-Kłosowicz et al. 2011; Jabłonka et al., 2012). PiDs were considered a major factor inducing the expression of COX-2 (Koistinaho and Chan, 2000; Miettinen et al., 1997). PiDs are caused by uncontrolled glutamate and potassium release in the ischemic tissue with prolonged elevation of extracellular K^+^ concentration and impaired energy-requiring K^+^ buffering by the astrocytic syncytium (Nedergaard and Hansen 1988; Hossmann, 1996; Dienel and Hertz, 2005). The extracellular K^+^ seems to be the main trigger for depolarization and for the accompanying depression of spontaneous activity (Hartings et al., 2017). The same mechanisms lead to SDs which are responsible for migraine aura in intact brains and for the sustained protein expression changes in humans after migraine (Charles and Baca 2013). SDs, which are experimentally induced by direct K^+^ application on the dura surface increase the concentration of COX-2 and MMP proteins and lead to wide-spread gene expression changes in the target hemisphere (Urbach et al. 2006). Since COX-2 post-stroke synthesis is mainly induced by the PiDs (Koistinaho and Chan 2000), we decided to check if SDs alone can affect plasticity that is induced by partial whiskers deprivation. In order to better understand metaplasticity changes after stroke, we also looked for other differences between the SDs induced by epidural KCl application and PiDs observed after cortical PtS.

Since post-stroke and post-SDs changes are characterized by similar dynamics of proteins changes (Gursoy-Ozdemir et al., 2004; Jabłonka et al., 2012; Urbach et al. 2006) and since some of the proteins were found in dendrites’ spines during synapse-remodeling and are therefore probably involved in cortical plasticity mechanisms (Ethell and Ethell, 2007; Fujioka et al., 2012), we decided to examine the presence of MMP-2, −3, −9, and COX-2 after SDs and PiDs. This is because these proteins seem to participate in the restoration of deprivation-induced plasticity in BF after PtS following intraperitoneal injections of FN-439 (MMPs inhibitor) or ibuprofen (unspecific COXs inhibitor and PPARs activator; Cybulska-Kłosowicz, 2011; Jabłonka et al., 2009, 2012). These studies showed that SDs induce additional COX-2 synthesis in the whole hemisphere at least up to the 5th day (Dihné et al., 2002; Horiguchi et al., 2006; Nedergaard and Hansen, 1988; Urbach et al., 2006), and MMP-9 overexpression was also demonstrated a few days after SDs (Gursoy-Ozdemir et al., 2004) as did PtS (Cybulska-Kłosowicz, 2011; Jabłonka et al., 2009, 2012). We therefor tested if protein distribution may be involved in post-stroke ExDP reduction (Cybulska-Kłosowicz et al. 2011; Jabłonka et al., 2007; 2012; Meighan et al., 2006; Nagel et al., 2005; Romanic et al., 1998; Urbach et al. 2006; Wang et al., 2008). We found that neither PiDs nor MMP-9, or −3, −2 or COX-2 proteins directly caused cortical post-stroke metaplasticity changes. A greater imbalance in interhemispheric activity equilibrium after stroke was the only difference between the effects of tested aspects of non-stroke and stroke depolarizations.

## Material and Methods

### Animals

The experimental protocols were in accordance with the European Communities Council Directive (86/609/EEC) and were approved by the local authority (Thueringer Landesamt, Bad Langensalza, Germany (02-20/05) and 1st Ethical Commission in Warsaw, Poland (270/2012, 790/2016).

Male Wistar rats weighing 290±20g (about 3 months old) were used for the experiments. Animals were kept in constant 12 hour light/dark conditions with food and water access *ad libitum*. Rats were randomly assigned to 15 experimental groups.

**We compared** (See Table1):

(**I**) whiskers cortical representations and
(**II**) general metabolic activity pattern

**between groups of:**

a. intact animals (n=6),
b. animals after ExDP induction, followed by cutting of all the whiskers except row B, for one month (**C**, n=6)
c. SDs induced by hyperosmolar KCl dura application (**SD**, n=8),
d. hyperosmolar NaCl instead of KCl application in sham animals (**SDS**, n=5),
e. PiDs induced by focal cortical stroke (**Pt**, n=6).

**Supp. Tab. 1:**
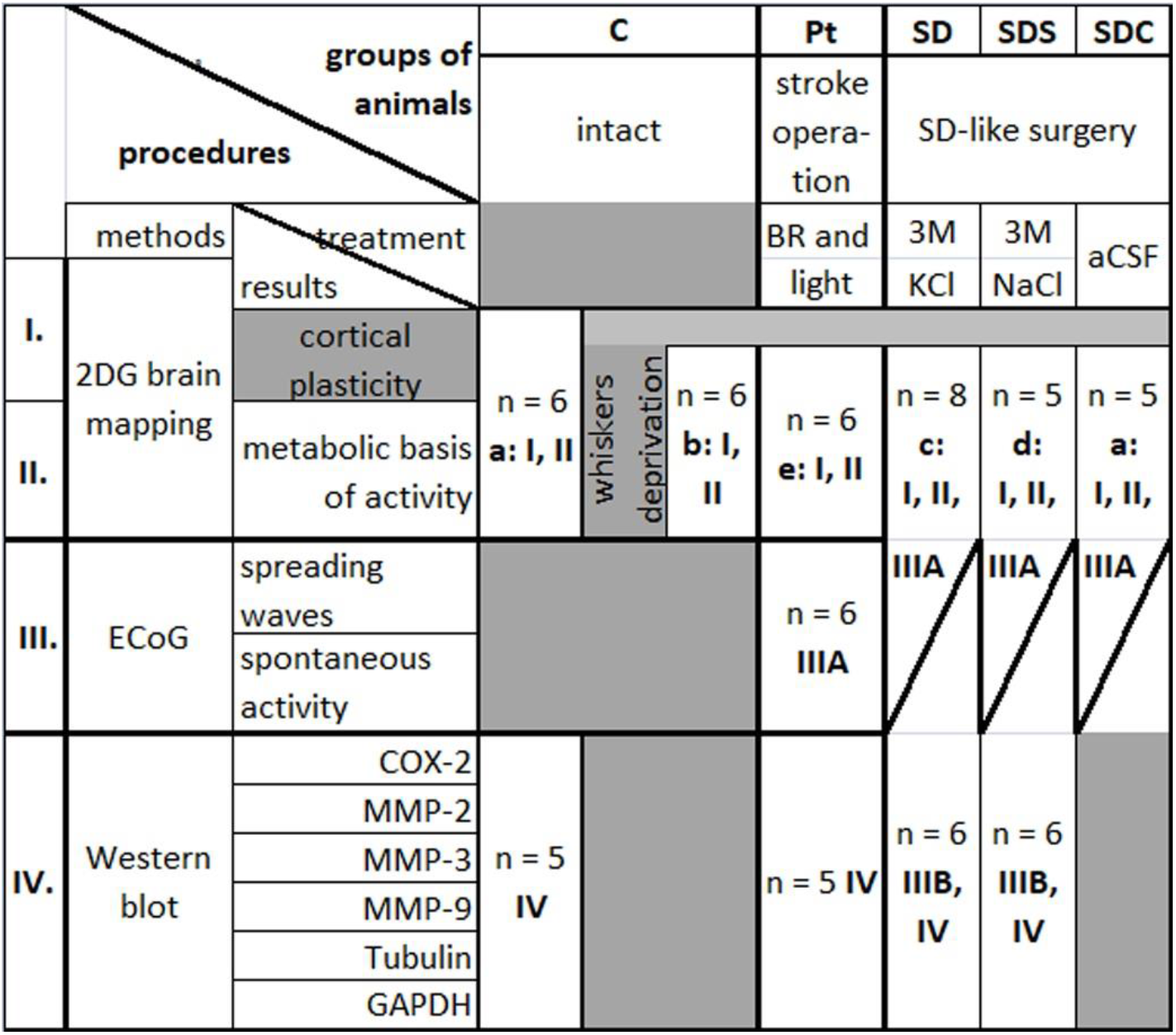
Groups of animals undergoing different treatments. **(C, Pt, SD, SDS, SDC),** and tested for changes in I-cortical plasticity, II-metabolic basis of activity, III-electrophysiological basis of activity and spreading waves characteristics, IV-protein changes.

**This was followed by comparing:**

(**III**) Electrophysiological features (n=6 for each group) and
(**IV**) Protein differences among the groups (n>5 in each group).

(The numbers from **I** to **IV** will be used as a reference to the specific experimental schemes related to these goals throughout the text).

#### Surgeries

During all surgical procedures, the animal’s heart and breathing rates were monitored continuously. The blood saturation with body temperature was also controlled (SOx 98% 1%; 36.5° **C±**0.5°C; MouseOx Plus, Starr). To prevent dehydration and maintain an adequate level of nutrition, at every second hour of narcosis the animals were given additional subcutaneous injections of 2 ml saline or Duphalyte (solution of amino acids, mineral ions, and carbohydrates; Pfizer OLOT, Spain).

##### SD-like surgery (Ib,Ic,II, IIIa, IIIb, IV)

30 animals had SD-like surgeries. (I, II). For ExDP studies, animals (n=18) were anesthetized with 2.5% isoflurane in N2O/O2 (60/30 l/h) and fixed in a stereotactic frame. The skull was exposed by an incision along the midline and two trepanations were drilled over the right hemisphere, (1.8 mm in diameter; coordinates: bregma −6.8 and bregma +2, both 2 mm from midline) leaving the dura intact. After reduction of anesthesia to 1.5% isoflurane, solutions were applied to the brain at the posterior trepanation (sol) for about 2 hours: 3M KCl (**SD** group), 3M NaCl (sham – **SDS**) or aCSF (control – **SDC**). Trepanations and wounds were then closed. To verify the occurrence of **SD** we concurrently recorded the electrocorticogram and direct current potential at the anterior position with a single electrode, by placing a glass electrode filled with aCSF (120 NaCl, 2 CaCl2, 5 KCl, 1.8 MgCl2, 10 HEPES, 1.25 NaH2PO4 and 10 Glucose: in mmol/L) and containing an Ag/AgCl wire on the dura exposed in the anterior craniotomy. An Ag/AgCl-reference electrode was placed subcutaneously in the neck.

During the operations intended for further studies of protein expression changes and for setting a more precise electrophysiological recordings the frontal trepanation was made slightly wider and a row of 16 electrodes (1.6 mm wide) were positioned on the dura to visualize direct spread of **SD** waves (coordinates: bregma +1—+4, 2,5 mm wide, for electrode positioning 5 mm from the midline, n=17) (Fig. 1A).

**Fig. 1.:**
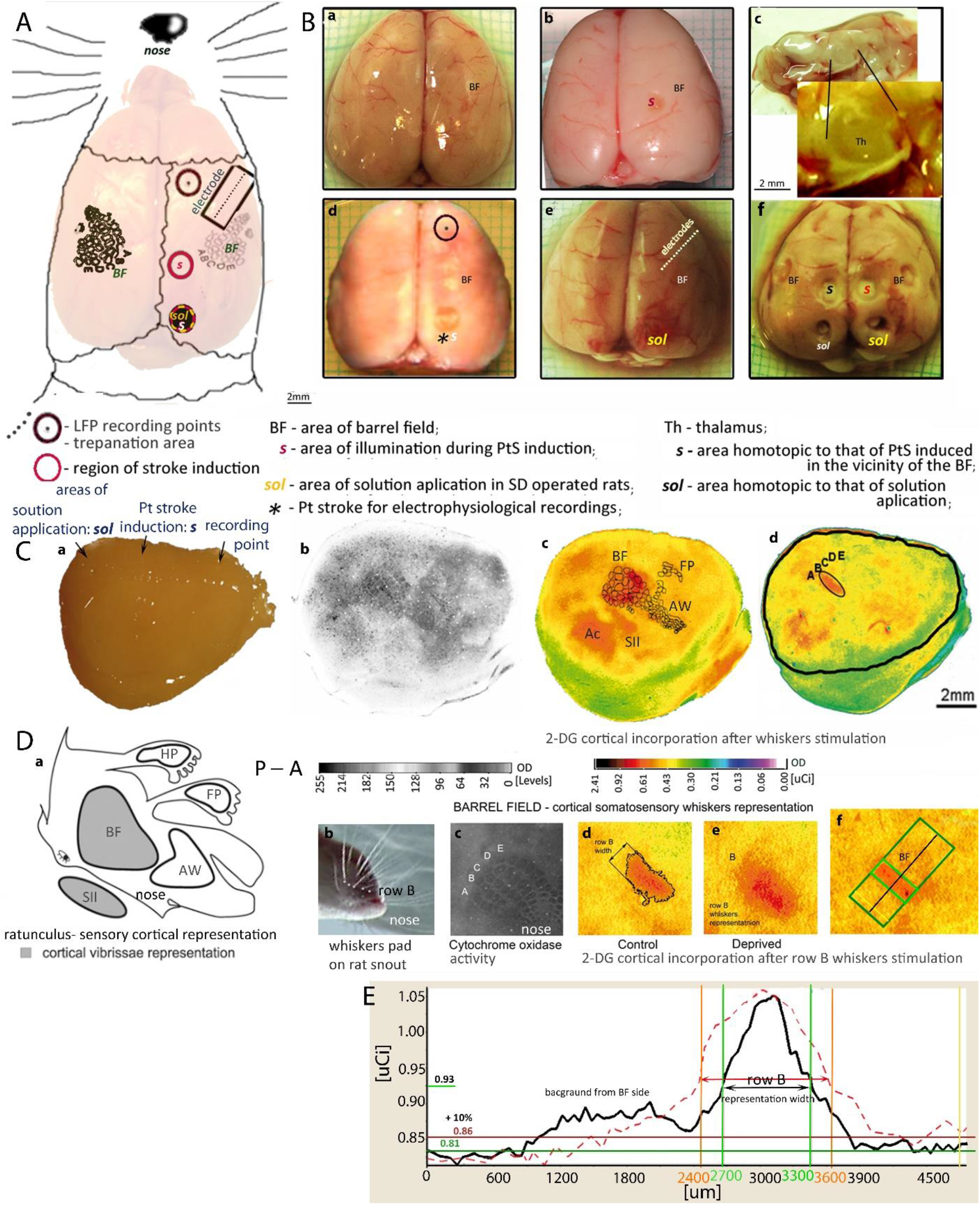
Examples of experimentally treated brains and further cortical treatment for activity mapping. **1A:** The schematic drawing shows foci of spreading depolarizations accompanying ExDP initiation (region of the photothrombotic stroke induction (***s***), solution application (***sol***)), location of the electrodes (the ECoG recording points) and skull trepanation areas (relative to the area of the barrel field, BF). **1B:** Isolated brains showing: 2DG brain mapping of ExDP (1B: a,b,d);electrophysiological recordings of PiDs and SDs (1B: c, d) and samples isolation for protein analysis (1B: a,b,d, e,f); from: Control (1B.a), **Pt** (1B: b,e,f) and **SD**/**SDS**/**SDC** (1B: e,f) animals. Brain prepared for thalamus (1B.f) and cortical samples (1B.e) isolation from BF, ***sol*** and ***s***. **1C:** Flatted hemisphere for tangential sectioning of the cortex showing 2DG brain mapping visualization (1C.a) with a marked position of ***sol***, Pt stroke induction (***s***) and ECoG recording point. Localization of the rats’ cortical whiskers somatosensory representations on a tangential section of layer IV (1C: b,c,d): stained by Nissl staining (1C.b) with 2DG-incorporation visualized on autoradiograms (1C: c,d), shown in pseudocolors. (AW) anterior whiskers representation; (FP) front paw representation; (SII) secondary somatosensory cortex. Measurements were also taken in (Ac) auditory cortex **1D:** A sketch of cortical vibrissal representation in SI and SII. (1D.a) “Rattunculus” with areas of increased activation during all the whiskers stimulation. The whiskers stimulation in intact rats activates contralateral BF and SII and do not activate anterior whiskers (AW), hind paw (HP) and front paw (FP) cortical representations. (1D.b) Photo of the whiskers pad on the rat snout showing row B. (1D.c) Magnification of the barrel field on the cortical tangential section after staining for cytochrome oxidase activity. (1D: d,c) a magnification of row B cortical representation, on digitized autoradiograms of tangential sections, in the control (1D.d) and deprived (1D.e) animals. 1D.f) Row B representation in the barrel field with the marked area of OD measure. **1E:** Comparison of the OD graphs showing 2DG incorporation in non-deprived (black line) and deprived BF (dashed red line). OD analysis by the MCID program, where the area exceeding the background over 10 % were treated as the activated representation of the rows B. The background mean OD was treated as a reference (right side of the graph). OD—optical density in arbitrary units with a direct set of 255 gray scale levels or calibrated to μCi radiation units of ^14^C-2DG; P–A—posterior to anterior (Supp. Fig. 5).

##### Photothrombotic stroke surgery (Id, II, IIIa, IV, V)

Photothrombotic stroke was conducted in the right hemisphere cortex of 17 animals. Animals were anesthetized and the skull was exposed as was described for SD-operated groups. Rose Bengal (0.4 ml, 10 mg/ml; 330000; Sigma-Aldrich, St Louis, MO, USA) was injected. After drilling the trepanation for electrophysiological recording the fibre-optic bundle light center (aperture=1.5 mm) was placed on the cleaned skull bone:

- (I, II, IIIa) over the back frontal trunk sensory representation (***s***, coordinates: bregma - 4.5 mm and 4 mm lateral from the midline (Fig. 1A)) in the vicinity of the somatosensory cortical representation of the whiskers where plasticity was induced;
- (IIIa) over the visual area of the occipital cortex to facilitate direct comparison of **SD** electrophysiological features to the PiDs occurring during ischemia (coordinates: bregma −5, 3 mm lateral (Fig.1A)) with corticogram measured from the dura above the prefrontal cortex by a single glass electrode (like during the described above SD-like surgery I, II; coordinates: bregma +2mm, 2 mm lateral to the right; with trepanation 1.8 mm in diameter).

##### Electrophysiological recordings (IIIa, IIIb)

The electrocorticograms (ECoGs) were recorded over a period of about 2h starting at least 2 minutes before the application of salt solutions in SD-operated rats or before Bengal Rose injection in **Pt** group.

The recording single glass electrodes (2 to 4MΩ) and/or sixteen, gold covered, tungsten electrodes (120 to 200MΩ) were placed on the dura surface as described in the section describing the surgery procedures. All data were transferred to a PC computer for online display analysis, and data storage using a high impedance amplifier (EXT-08, NPI, Tamm, Germany (for glass electrodes) or for the tungsten electrodes the A-M Systems amplifier, model: 3600 with preamplifier attached directly to the socket of the electrodes holder). Signals from all electrodes were band-pass filtered (0.3 – 5000 Hz) and transformed by CED 1401 Plus analog/digital converter. Software Spike2 (Cambridge Electronic Design, UK) were used for signals digitalization at 19 kHz sampling rate per channel.

### Whisker deprivation and ^14^C-2-DG brain mapping

#### Deprivation and brain tissue preparation (I)

Immediately after the surgery, all left-side whiskers (contralateral to the SD) except that of row B were trimmed close to the skin, with trimming repeated every second day. After four weeks of deprivation, we performed ^14^C-2-deoxy-D-glucose (American Radiolabeled Chemicals; St. Louis, MO, USA) brain mapping. Rats were immobilized and whiskers were cut bilaterally except rows B. The 2-DG was injected intramuscularly (7 μCi/100g body mass), and rows B of whiskers on both sides of the snout were stimulated simultaneously at a frequency of 2 Hz. After 30 minutes of stimulation, animals were deeply anesthetized by intraperitoneal injection of 1 ml Vetbutal and intracardially perfused with 4% paraformaldehyde (Sigma-Adrich; St.Louis, MO, USA). The brains were removed and cortices of separated hemispheres were flattened between glass slides to 3 mm, snap frozen in heptane at - 70°C and stored at −80°C.

#### 2-DG incorporation imaging (I, II)

Flattened hemispheres were cut tangentially on a cryostat at −16°**C** into 40 μm sections, which were collected alternately on slides and coverslips. The sections on coverslips were immediately dried and exposed on X-ray film (MIN-R 2000; Kodak) for 2 weeks together with a set of [^14^C] standards (ARC 146; American Radiolabeled Chemicals; St.Louis, MO, USA;) and later stained in Nissl staining visualizing neuronal polyribosomes. The remaining sections were stained for mitochondrial cytochrome oxidase activity to identify the barrel field and the lesion regions.

### Protein extraction and Western blot (IV)

#### Tissue samples

The concentrations of proteins (COX-2, MMP-9, MMP-3, MMP-2) were analyzed using Western blot analysis for homogenates of chosen areas isolated 24 h after SDs episodes or stroke induction. Tubulin, GAPDH and total protein with Coomassie Blue were used as a loading control. Tissue samples were collected from rats killed by decapitation. Brains were dissected on ice and tissue samples were immediately transferred to proteinases inhibitors (10% PiC solution), frozen in the liquid nitrogen and stored at −70°C.

The cortical samples were isolated by an absorption (with 2 mm in diameter tube) of the column from the cortical surface to white matter (white matter excluded). The tissue samples were taken bilaterally from the barrel fields (Strominger and Woolsey, 1987), stroke area (***s***) and area of solutions application (***sol***) and additionally from the auditory, prefrontal cortex and thalamus (Katagiri et al., 2010).

#### Proteins isolation

After the addition of an ice-cold RIPA buffer (50 mM Tris, pH 7.5, 150 mM NaCl, 1% NP40, 0.5% sodium deoxycholate, 0.1% sodium dodecyl sulfate, 1 mM ethylenediaminetetraacetic acid, 1 mM NaF, Complete Protease Cocktail and Phosphatase Inhibitor Cocktail 2) the tissue samples were homogenized by sonication, centrifuged, and stored at −80°C.

#### Western blots

Proteins were separated in sodium dodecyl sulfate-acrylamide gels and transferred to PVDF membrane. PageRuler™. Prestained Protein Ladder was used as a size standard (26616, Thermo Fisher Scientific). The membranes were blocked with non-fat dry milk and incubated overnight with primary antibodies at 4°**C** (Abcam polyclonal rabbit IgG: Anti-MMP2 – ab37150, Anti-MMP3 – ab53015, Anti-MMP9 – ab38898, Anti-COX-2 – aDb15191 and Abcam mouse monoclonal anti-beta III Tubulin IgG2a – ab78078, anti-GAPDH, SantaCruz, SC-25772). After washing, the membranes were incubated with peroxidase-coupled secondary IgG antibodies (anti-Rabbit–A0545 or anti-Mouse –A9044, Sigma Aldrich) for 2 h at room temperature. Staining was then visualized by chemiluminescence. The images were captured using an ImageQuant LAS 4000.

### Data analysis and statistics (Dast)

#### DaSt: 2-DG incorporation, histological stainings and Western blots

The autoradiograms, blots and histological images were analyzed by a computer image analysis program (MCID; InterFocus Ltd., Cambridge, UK), which measure the optical density (OD) with 8-16 or 24 bit accuracy. The OD and distance were calibrated. Blots densitometric analyses were performed using Quantity One 1-D software (Bio-Rad) and compared with that made by MCID.

The differences between groups were tested using a multi-factor ANOVA and a post-hoc Tuckey test (for unequal N). The significance of the differences between the hemispheres was calculated with a two-tailed paired Student’s T-test.

#### DaSt: Histological tissue analysis (I, II, IIIa)

The color images of histologically stained sections were coded in 24 bits. The software allowed us to display on screen the image of a Nissl stained section from which the autoradiogram was obtained and adjacent sections stained for cytochrome oxidase activity, and to superimpose these images on each other so that the relation of the barrel field to the position of the electrodes, lesions or solution application areas could be precisely measured.

The tangential sections of KCl and NaCl treated groups do not permit a closer study of the lesion size and distribution. Therefore, we measured the lesion diameter on the surface of the brain (the widest place) and its shortest distance to the barrel cortex on the surface of the unprepared flattened hemisphere and on slices from layer IV, where the barrel field was visible. The diameter of the stroke was also measured on layer IV of the tangential sections. Lesions depth was measured based on the extrapolated edge of the tangential slice in the location of the tissue destroyed by the lesion, and its distance to the deepest place of lesion bottom (Fig. 1A).

#### DaSt: 2-DG incorporation (I, II)

The 2-DG incorporation level was measured on standardized images of the sections autoradiograms. The grayscale images of autoradiograms were analyzed with 8 bits accuracy with 255 gray scale levels.

#### DaSt: Experience dependent plasticity (I)

The width of the labeled cortical representation of row B whiskers was measured in layer IV. The pixels with 2-DG-uptake intensity that were consistently above 15% of the mean surrounding cortex incorporation were considered as the labeled representation. The results were averaged for all sections from layer IV of one hemisphere.

#### DaSt: Metabolic correlates of cortical activity (II)

Metabolic rate in different cortical regions was based on densitometry measures from [^14^C] 2DG autoradiograph. After establishing the optical density of the autoradiograms within the range of [^14^ C] standards and ensuring that the standards optical density increased in a linear manner, autoradiograms were compared with stained brain sections. On the assumption that the OD in autoradiographs is proportional to the concentration of the [^14^C] isotope that in turn reflects local glucose consumption (Sokoloff et al. 1977), we converted the OD values from digitized images into μCi based on the ODs of the [^14^C] standards that were co-exposed with the cortex sections. [^14^C] 2DG uptake was measured on tangential sections and averaged for cortical layers II/III, IV and V/VI. The degree of shift in the balance of activation between the treated and “intact” hemispheres was characterized by the laterality index. Regions of interest were defined on Nissl stains and stained for cytochrome oxidase activity.

#### DaSt: Western blots (IV)

Since we used 8 bits grayscale images, 255 gray scale levels were chosen for 2DG incorporation analysis (Fig. 1D).

After background subtraction, the protein of interest was normalized relative to housekeeping proteins and multiplied by the surface of the analyzed protein strip (mostly ß-tubulin but for ***s*** tissue samples also GABDEH was used due to evident inter-group differences in tubulin concentration). The normalized number of the protein amount [An] was used for data presentation. When the OD exceeded the background by 10 % (5 An) (Fig. 1D & Supp. Fig. 2) protein presence in homogenate was assumed. The final An for each animal were compared between groups present on one blot; inter-blot comparisons were also made.

The correlations for proteins concentration in the different groups, in different brain areas, and between particular animals in each group, were checked and compared for each blot. When a band of one or two rats enhanced the SD above the mean value they were mentioned separately (as “+ xx [An]”).

#### DaSt: Additional bands specific for groups

Since we found that some protein bands of untested functional significance were reliably correlated with some of the experimental groups (Supp. Fig 3), we measured their OD and used them as an additional source of information (Supp. Fig. 5 and supplementary data). Heavier (protein+X) and lighter (protein-X) bands were considered separately. Comparison of groups with unequal variances in the additional bands was performed by using the Mann-Whitney (MW) test, which was supplemented with Bonferroni correction for testing six hypotheses. Thus, the established statistically significant p-value was p=0.008.

#### DaSt: Electrophysiological brain activity (III)

Analyses of the electrophysiological data were made using Python programing language and Matlab (MathWorks) statistical tools.

### DaSt: Electrophysiological data pre-processing

#### DaSt: Single electrode recordings (IIIa)

To visualize basic electrical activity during spreading depolarization we recorded raw signal from one glass electrode at the dura level. To establish depolarization time, we applied a “moving variation” method for low pass <0.1 Hz filtered signal. Depolarization was averaged across depolarization trials in PiDs (n=16) and SDs (n=10) (Fig1A) and then compared between groups (Fig 1B).

We filtered signal 0.1-100 Hz with a third order Butterworth filter. We computed spectrograms for 50 minutes band-pass filter signal. In order to search for electrophysiological differences between PiDs and SDs, we have chosen fragments of ±250 seconds from continuous signal around the trough of the significant depolarization sites and we treated them as trials for further analysis. We then filtered signal 0.1-50 Hz and calculated spectrograms using 5-sec non-overlapping window.

#### DaSt: Multielectrode recordings (IIIb)

The SDs signal was recorded with 14 channel tungsten electrode with 19.6 kHz frequency. We filtered the signal with a 4th order filter, low-pass from 1 Hz. The sampling rate was reduced from the original sampling rate by a factor of two, from the 19608 Hz to 9804 Hz.

#### DaSt: Electrophysiological statistics (IIIa, IIIb)

Most differences between the frequency of the bands were tested using a multi-factor ANOVA and a post-hoc Wilcoxon test. To estimate significant differences between groups we used the resampling method. We picked (10,000 times) random depolarization fragments and calculated mean spectrograms for these new random sets. As a statistic, we used the difference in every “pixel” calculated for groups and compared it with the original difference between PiDs and SDs. This provided us with a p-value for every “pixel” for average spectrogram picture (Supp. Fig. 1).

### CSD analysis

To estimate the distribution of current source and sinks of the depolarization event recorded at multiple sites, we used kCSD reconstruction method (Potworowski et al., 2012). We assumed constant and isotropic tissue conductivity for CSD reconstruction.

## Nomenclature

For easier accommodation of the data and easier document navigation, we use abbreviations and different letter formats. Brain regions examined in the experiment are abbreviated in bold and italicized (***BF, sol, s, Th***); Experimental groups are abbreviated and denoted by bold uppercase letters (**C, SDC, SD, SDS, Pt**). Each group is marked by corresponding color on each figure (**C** – green, **SDS** – blue, **SD** – yellow, **Pt** – red)

## Abbreviations

2-DG: ^14^C-2-deoxyglucose
Au: auditory cortex
BF: barrel field
ECoG: electrocorticograms
ExDP: experiencedependent plasticity
Hp: hind paw
*MW*: Mann-Whitney test
OD: optical density
OS: open source
PiD: peryinfarct depolarization
PtS: photothrombotic stroke
*s*: stroke area with the PtS induced in the **Pt** group
SII: secondary somatosensory cortex
*sol*: solution application area
SD: spreading depolarization in contrast to experimental group **SD** and *SD* – standard deviations
*th*: thalamus

## Results

### Histopathological verification (I, II, IIIA)

Both the experimental and sham animals had a surface cortical lesion in the brain location where the solution was applied and PtS induced.

#### After SD-like surgery

Lesions induced by high osmolarity solution were 300 um±120 um in depth and diameter on layer IV: for NaCl – 1±0.5 mm and for KCl – 1.5±0.2 mm. The diameter of the cortical surface decompositions in layers II/III were 3.4 ±0.3 mm for NaCl, and 4.3 ±0.9 mm for KCl. They were localized in the secondary visual cortex (V2M) and concentric surroundings: medially, the retrosplenial agranular cortex (RS) and laterally, the medial primary monocular area (V1M, Urbach et al. 2006, Fig. 1A).

The high osmolarity-induced lesion edge was in a distance of 4 mm±0.4 mm from the B1 barrel (Fig. 4). There were no structural differences in the distance from 0.2±0.1 mm around the edge of the ***sol*** area, nor in remote areas. The group after a CSF application did not have the lesion on the cortical surface showing only a small oedema which was not statistically significant when compared to intact brain deformations (without craniotomy).

**Fig. 2.:**
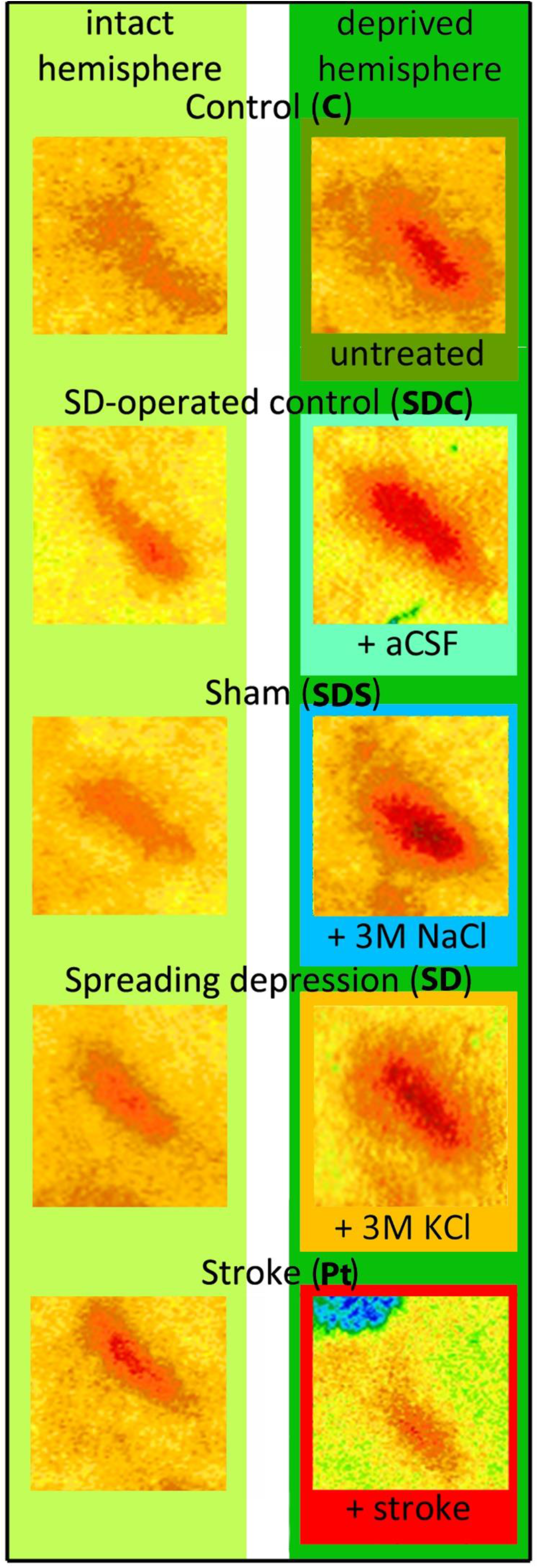
**Rows B cortical representations** in the deprived and non-deprived hemispheres: in control animals (C), after PtS (Pt group); after cortical treatment with: aCSF (to SDC animals) or application of the highly osmotic solutions of NaCl (in SDS) or KCl (in SD) to V2 cortical surface. The magnified parts of autoradiograms of cortical tangential sections, visualized by the exposition of the brain slices to ^14^ C-2DG incorporations during bilateral, simultaneous rows B whiskers stimulation (Fig.1.E.).

**Fig. 3.:**
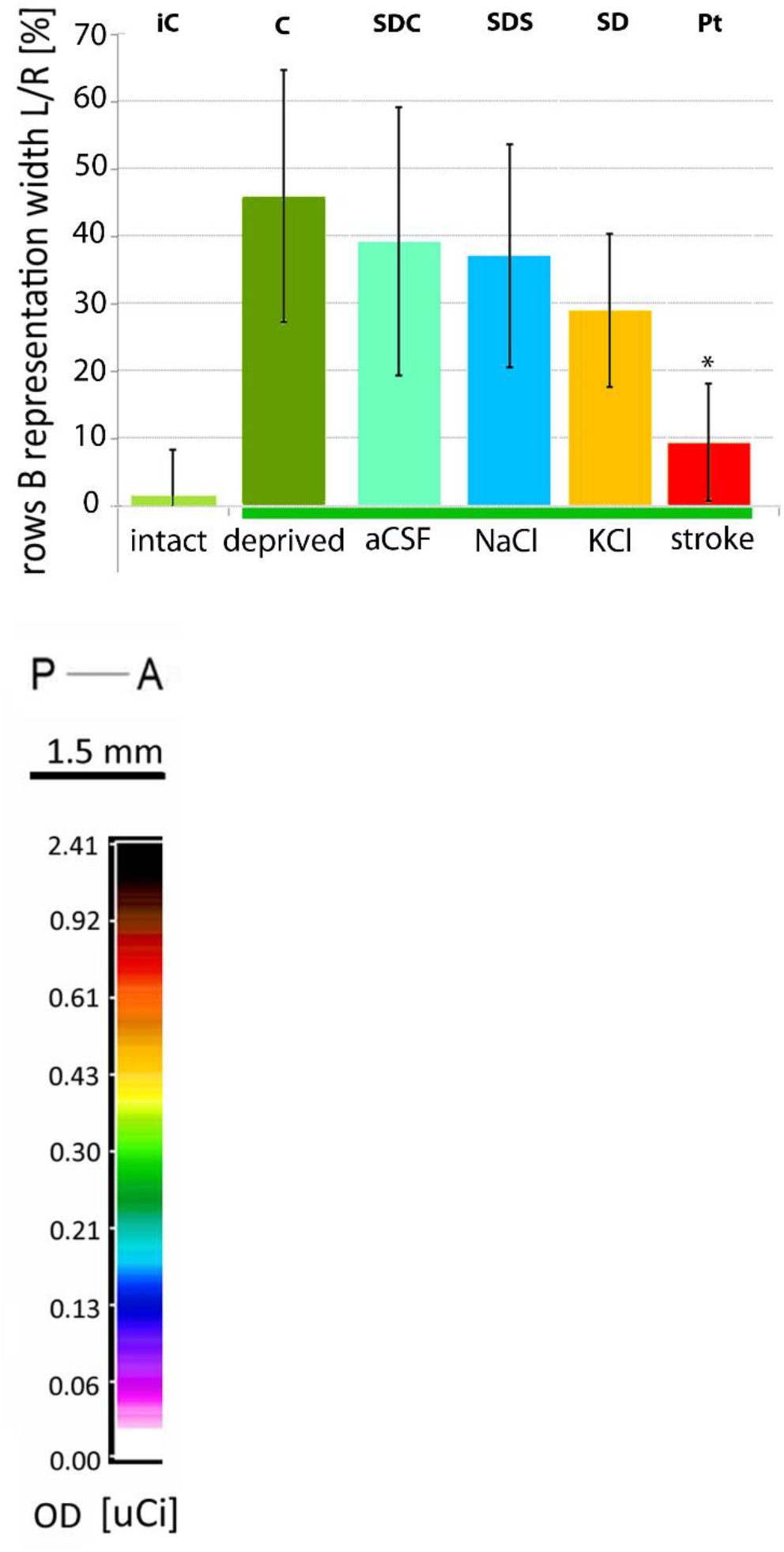
The ratio (in percentage) of row B representations width in the non-deprived and deprived hemispheres. There was a significant difference in each experimental group (except the Pt). The Pt-group had significantly lower width in comparison with every other group (p < 0.001).

**Fig. 4.**
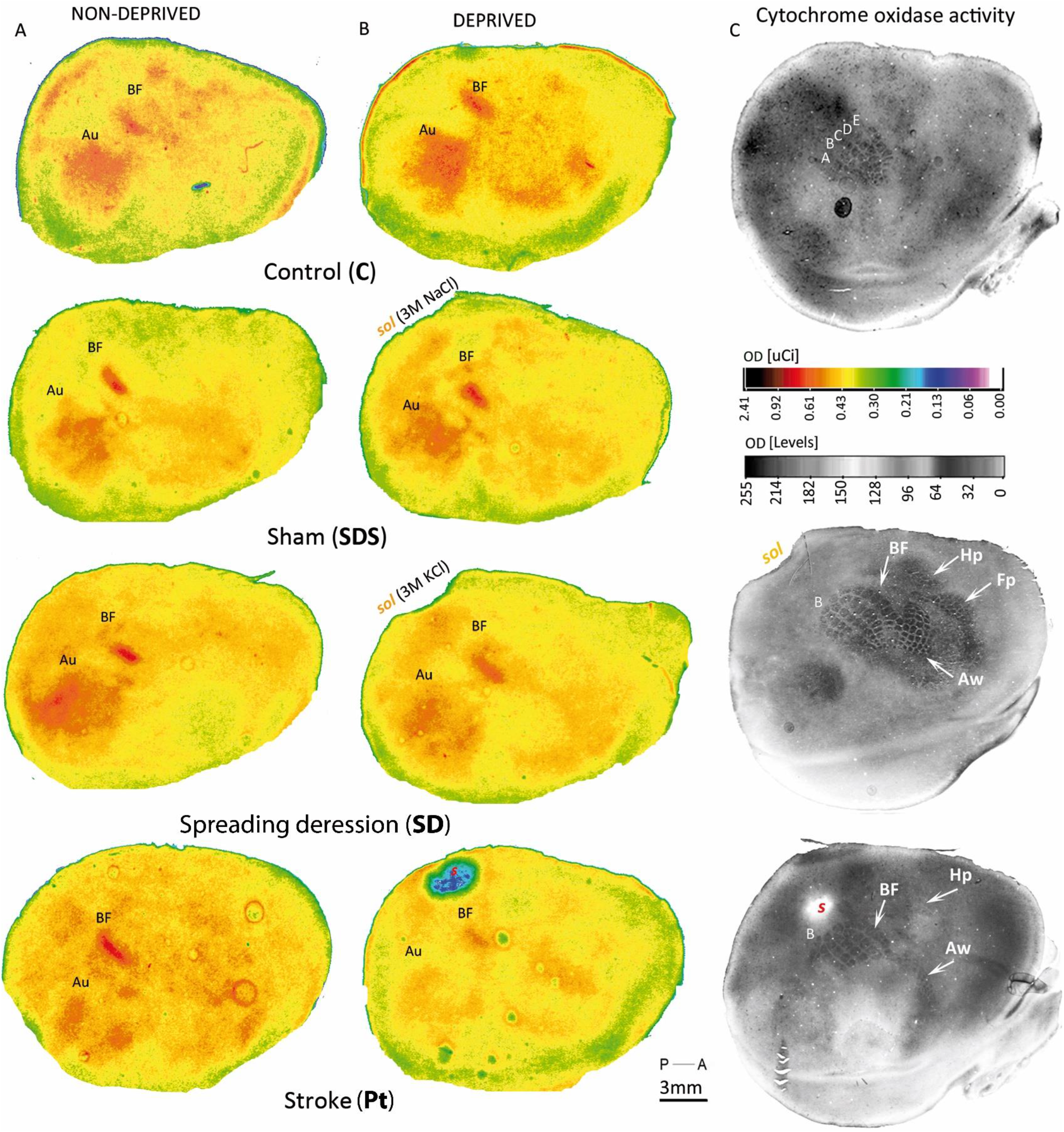
**Cortical metabolic activity during brain mapping following bilateral whiskers stimulation of row B** one month after treatment. 2DG incorporation shown on autoradiograms of the entire hemispheres, of layer IV, and of tangential sections. Comparison between the experimental groups (control (**C**), and SD-operated group: sham (**SDS**) and after spreading depression **SD** and stroke (**Pt**) with intact (left, A) and deprived (right, B) hemispheres (for more details see Table 1). On the right, the equivalent sections stained for cytochrome oxidase activity (C). A and B are shown in pseudocolors (OD—optical density in arbitrary units).

#### After PtS induction

PtS located in a part of the trunk sensory representation had the same diameter at the brain surface as that seen in the layer IV (1.5±0.5 mm) and was slightly smaller in the supragranular area of layers V, VI (1.3 mm±0.2 mm), leaving white matter intact. However, the CO staining showed a decreased level of optical density at a distance of up two 2 mm from the stroke core center (Fig. 4). The volume of the stroke core, where necrotic cells dominate, was 0.8 mm^3^±0.3 mm^3^. The lesion edge was at a distance of 1200±208 μm from the B1 barrel (Fig. 4). There were no structural differences in the distance 0.3 ±0.2 mm in the stroke core and remote areas of the cortex.

### 2-deoxyglucose (2-DG) brain mapping (I, II)

A multi-factor ANOVA analysis for all the groups revealed significant changes after PtS and SDs in the interhemispheric relation of 2**-**DG cortical incorporation (**II**, F_(2.89)_=5.38, p=0.006) and cortical row B representation area (**I**, F_(2.95)_=2.77, p=0.001), as well as in row B width (**I**, F_(2.09)_=10.34, p < 0.0001).

### Experience-dependent plasticity (I)

After one month of unilateral deprivation of all but one row of whiskers, we observed a widening of the non-deprived row cortical representations in all experimental and control groups (Fig. 2 & 3).

In animals with SDs and a focal lesion induced by 3M KCl, the difference between row B representations was the same as in the both **SDS** and **SDC** groups (p<0.05). The mean percentage difference between deprived and non-deprived sites was like that found in the intact control group **C** (42%±18%). In animals with focal PtS the difference was 11%±9%.

The width of the spared row B representation in the deprived hemisphere of the **Pt** animals differed significantly from corresponding areas in all the other groups with whiskers deprivation: **SD, C** (p<0.005) and **SDC** (p=0.04). There were no statistically significant differences between SD-operated groups (Tab. 2, Fig. 2, 3; Supp. 1).

**Tab. 2:** Non-deprived row B cortical representation width showing the percentage of interhemispheric difference and its significance; p<0.05

**Experience-dependent plasticity (I)**
| Groups | row B representation width<br>[μm] |  | interhemispheric<br>difference of row B<br>representation [%] | significance<br>of the<br>difference |
| --- | --- | --- | --- | --- |
|  | hemisphere |  |  |  |
|  | undeprived | deprived |  |  |
| C | 814 ± 15 | 1160 ± 121 | 42 ± 14 | p = 0.009 |
| SDC | 798 ± 108 | 1075 ± 132 | 35 ± 27 | p = 0.03 |
| SDS | 779 ± 123 | 1023 ± 192 | 33 ± 26 | p = 0.04 |
| SD | 781 ± 81 | 1036 ± 192 | 42 ± 18 | p = 0.002 |
| Pt | 821 ± 113 | 906 ± 76 | 11 ± 9 | p = 0.02 |

### Metabolic correlates of cortical activity (II)

Since in the poststroke brains, spontaneous activity seemed to have an important impact on cortical map reorganizations (Jabłonka et al. 2009), we examined the activity-related metabolic changes by examining the 2-DG incorporation level. There were no significant differences between the groups -? with regard to 2DG incorporation in the relation of 2-DG incorporation level of all measured areas was comparable between and within the groups and differed significantly from that in the **SD** group (p =0.004, Table 3b).

**Tab.3:** 2DG incorporation [μCi/g] in cortical rows B representations (A.) and in entire sections of the intact and treated hemispheres (B.); percentage of significant differences are shown; p<0.05.

| A<br>Groups | 2-DG incorporation in cortical rows B representations [$\mu\text{Ci/g}$] | | interhemispheric difference of 2DG incorporation in cortical row B representations | significance of the inter-hemispheric difference |
| --- | --- | --- | --- | --- |
|  | hemisphere |  |  |  |
|  | intact | treated |  |  |
| C | $1.08 \pm 0.1$ | $1.08 \pm 0.1$ | $1 \pm 0.1$ | |
| SDC | $1.03 \pm 0.2$ | $1.16 \pm 0.3$ | $1.05 \pm 0.07$ | |
| SDS | $1.28 \pm 0.6$ | $1.32 \pm 0.6$ | $1.04 \pm 0.1$ | |
| SD | $1.23 \pm 0.1$ | $1.13 \pm 0.2$ | $1.08 \pm 0.5$ | 8% |
| Pt | $1.17 \pm 0.2$ | $1.13 \pm 0.1$ | $0.96 \pm 0.04$ | |

| B<br>Groups | 2-DG incorporation in cortical rows B representations [$\mu\text{Ci/g}$] | | interhemispheric difference of entire sections 2DG incorporation | significance of the inter-hemispheric difference |
| --- | --- | --- | --- | --- |
|  | hemisphere |  |  |  |
|  | intact | treated |  |  |
| C | 1.02 ± 0.2 | 1.04 ± 0.2 | 1.02 ± 0.04 | |
| SDC | 0.99 ± 0.2 | 1.02 ± 0.3 | 1.03 ± 0.06 | |
| SDS | 1.09 ± 0.4 | 1.09 ± 0.5 | 1.00 ± 0.04 | |
| SD | 1.06 ± 0.2 | 1.02 ± 0.2 | 1.07 ± 0.09 | 7% |
| Pt | 1.19 ± 0.04 | 1.03 ± 0.1 | 1.13 ± 0.08 | 13% |

The interhemispheric differences of **SD** animals disappeared when we normalized the values relative to the auditory cortex, which was treated as an activated reference area (layer IV: **Au** – by 7 %, p=0.004, row B – 8%, p=0.007, layers V/VI: **Au** – by 5 %, p=0.05, row B – 4%; p=0.01); it had itself the interhemispheric shift by 7 % in layer IV (LH – 1.27±0.16 μCi/g, RH – 1.19±0.11 μCi/g).

One month after the stroke, the post-stroke group (**Pt**) had a shift ofinterhemispheric relation which was 13% higher in the intact hemisphere (Fig. 5, Tab. 3b) than in the treated one. Both the **Pt** and tested regions, but the relation between the hemispheres differed between the groups. No statistically significant interhemispheric imbalance were found in the **C, SDC,** and **SDS** groups.

**Fig. 5.:**
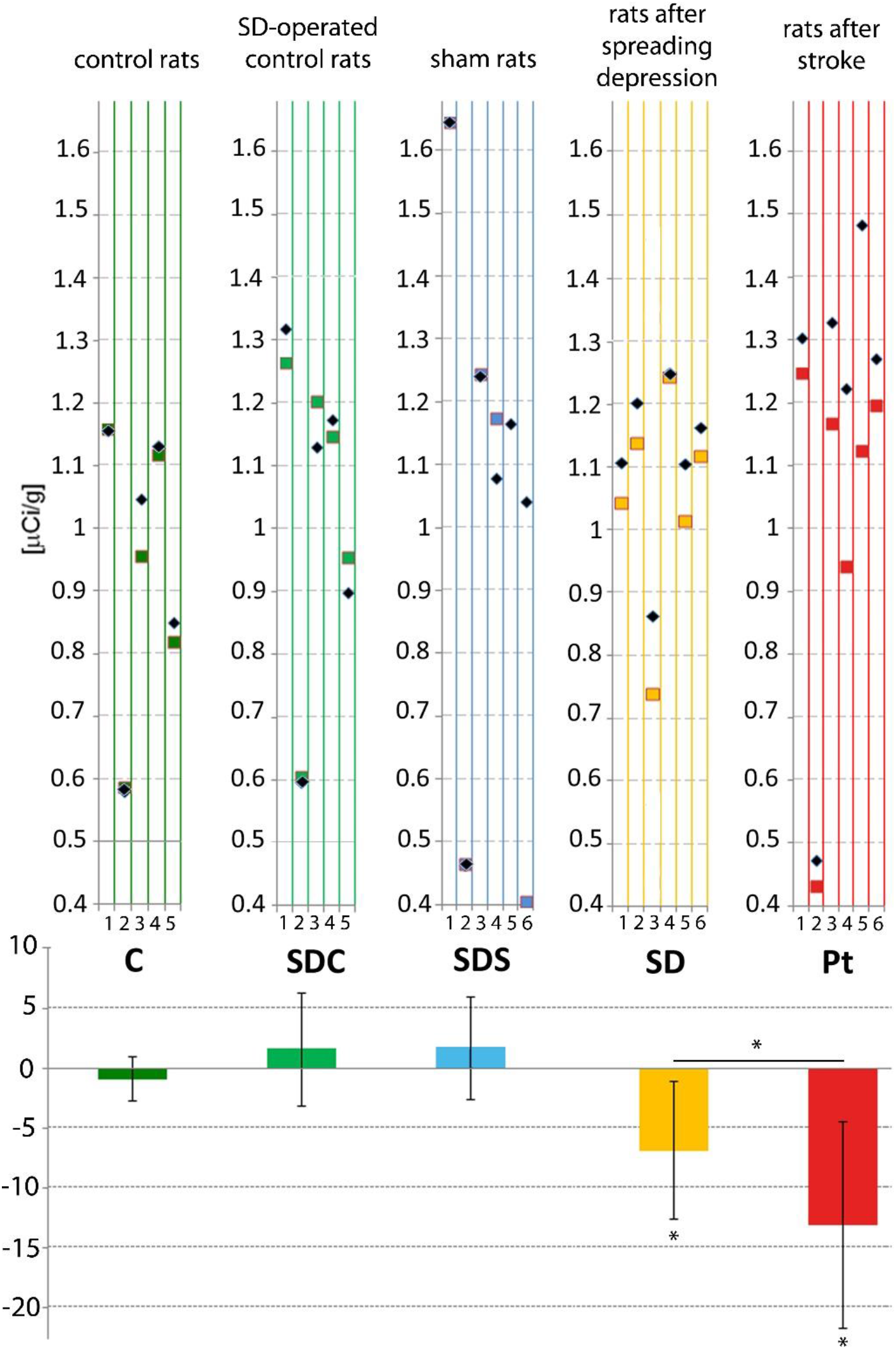
Cortical layer IV: 2DG incorporation in the treated and intact hemispheres. Upper graphs represent individual results for each tested animal. Squares represent right hemisphere, diamonds – left hemisphere. The lower figure shows the interhemispheric comparison of the mean ratio (right to left hemisphere) of 2DG incorporation in cortical layer IV presented as a percentage. According to ANOVA test, the 2DG incorporation was significantly lower - by 7% - in treated (right) hemispheres in **SD** group (p=0.006) and by 13% in the post-stroke **Pt** group (p=0.03). The laterality index was greater in **Pt** than in **SD** animals by 6 % (p=0.006)

In the control group (**C**), the interhemispheric ratio of 2DG incorporation in the representations of stimulated rows B was 1± 0.1 μCi/g. It was significantly higher than in surrounding non-stimulated cortex (0.89±0.04 μCi/g from the side of BF center — side of row **C** and 0.81±0.2 μCi/g laterally — from the side of row A, p > 0.0001) and this tendency did not differ between groups (Table 3a).

For all the measured areas in SD-operated groups, only the **SD** group had significantly lower 2-DG incorporation in layers IV and V/VI in the treated hemisphere, and 8% lower incorporation in the entire non-stimulated area of the treated hemisphere a month after the SDs episode. For the rest of SD-like operated rats, the interhemispheric**SD** groups differed significantly from controls (**SD** p=0.03, **Pt**: p=0.004).

These results allow us to combine groups **C, SDC,** and **SDS** into a single super-group with stabilized interhemispheric relation of activity, and to combine the **SD** and the **Pt** groups into super-group in which the interhemispheric activity equilibrium was affected. There were no differences within these two super-groups with respect to the interhemispheric ratio, but the difference ratio between the groups was pronounced (p=7E-05). However, when we compare the laterality index it was higher after stroke by 6% (p=0.009) than after SD (Fig. 5).

### Electrophysiological brain activity (III)

Looking for other differences between post-stroke and post-spreading depolarization conditions, we examined spontaneous cortical activity from electrocorticograms (ECoGs) recorded for 2h after the beginning of PtS induction (the light turned on) in the **Pt** group, and 2h since solution application in the **SD** group. The signals were compared to spontaneous activity in **SDC** animals after an aCSF cortical application and shams (**SDS**) with 3M NaCl application.

#### (IIIA)

The ECoG pattern of operated animals was normal, similar to isoflurane anesthesia in rats (Michelson and Kozai, 2018) in each group when monitoring started. The first depolarization appeared 42±23 [s] in **Pt** and 10±2 [s] in **SD** groups, after rose Bengal injection or KCl application respectively.

The significantly different was the duration of the depolarization wave of SDs 92±59 [s] and PiDs 108±39 [s] (F=4.53, p=0.03). Measurements of slow wave [<1Hz] oscillations showed no significant differences between **SD** and **Pt** groups. SDs and PiDs frequency (SDs: 3.8 ±1.9 Hz, PiDs: 2.9±1.3 Hz) and the amplitude from the baseline (SDs: 0.47±0.2 mV, PiDs: 0.48±0.5 mV) and the area below the depolarization wave were similar. The amplitude, however, was higher in PiDs when we compared it between the minimal and the maximal values within the analyzed period (p=0.042).

During depolarizations, a spreading depressions appeared — a pronounced silencing of the spontaneous oscillations over 3 Hz — in both **Pt** and **SD** animals (Fig. 6). The spectrograms did not reveal any difference in any frequency band between SDs and PiDs. In the **SD** group, the silencing of the oscillations exceeded the depolarization some time later (Fig. 7). However, no significant difference between PiDs and SDs were found.

**Fig. 6.**
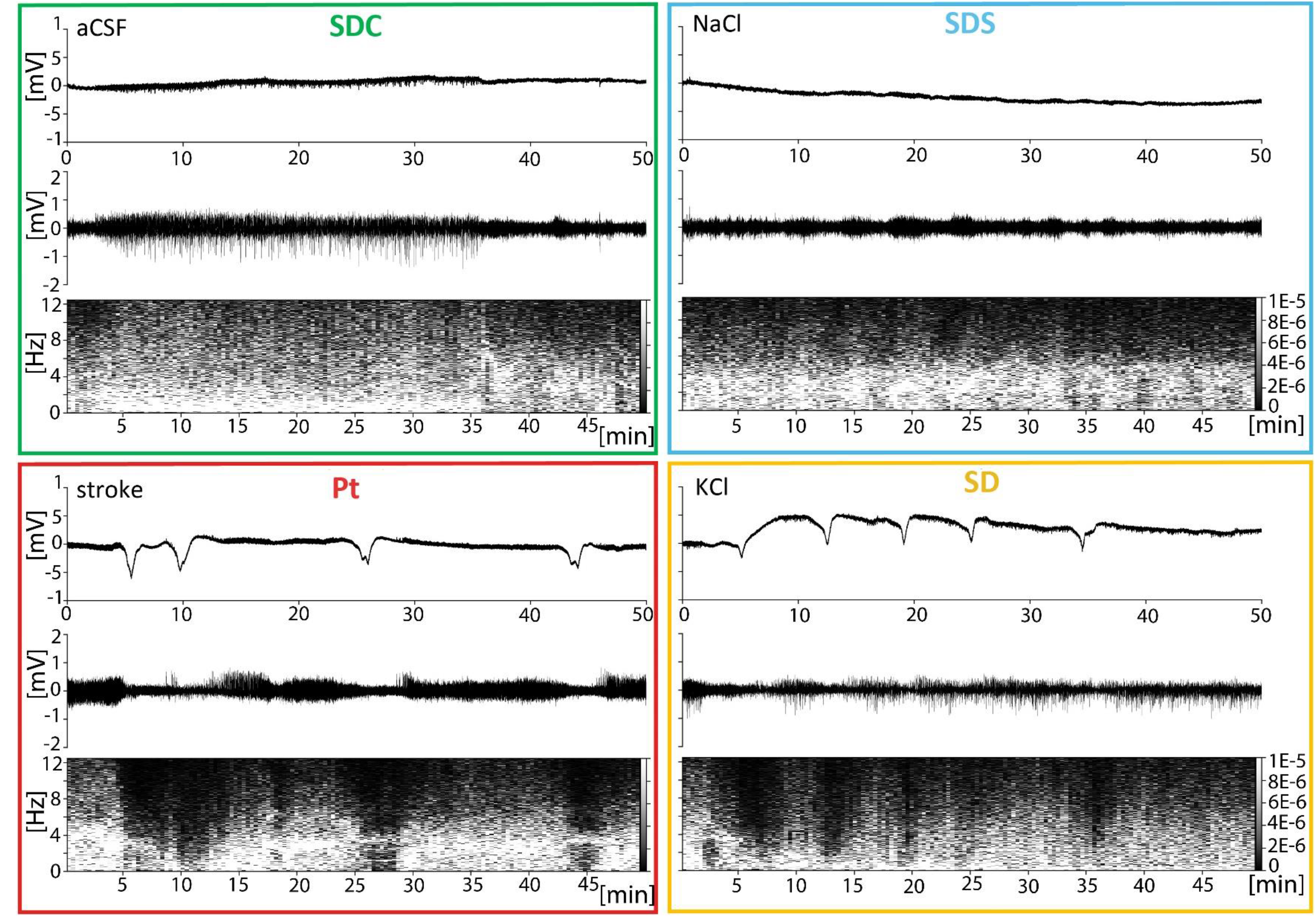
Plots of recordings for spreading depolarizations. Top pictures show raw ECoG signals. Signals in the middle pictures were filtered 0.1 – 100 Hz with Butterworth 3 rd order filter. Bottom panels present spectrograms computed from the filtered signal (See in methods - DaSt: Electrophysiological statistic). We observe a significant reduction of broad range (4-12 Hz) of ECoG oscillations that correlates with depolarization events both in Pt and SD experimental conditions. SDC (with aCSF) (A), SDS after 3M NaCl application (B), Pt during and after PtS (C) and 3M KCl in SD group (D), (Supp. Fig. 1, 2);

**Fig. 7.**
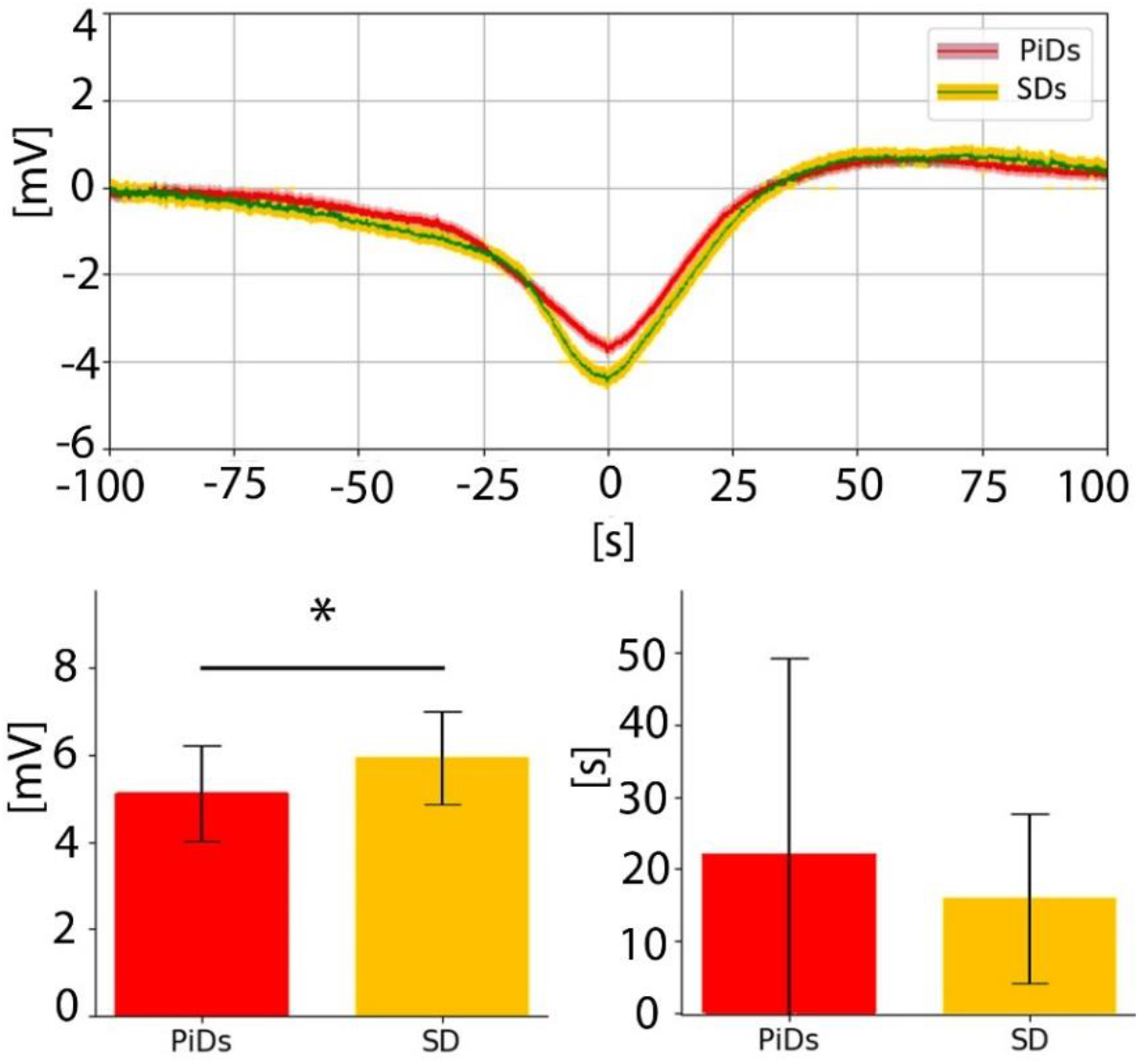
**Comparison of mean events of depolarizations:** evoked by KCl application (yellow line), poststroke PiDs (red line). Top panel presents the mean signal of detrended depolarizations (baseline translated to 0 amplitude). Bottom left: Amplitudes differences between PiDs and KCl. Bottom right: no difference in the time duration of spreading depolarization. The statistic was computed using the resampling method (Supp. Fig. 6-1).

#### (IIIB)

The multielectrode recordings in **SD** animals allowed us to visualize four types of depolarization waves traversing the space: 1) classical **SD-**like waves when depolarizations appeared on three or more adjacent electrodes in a radial epicentral direction from ***sol***, the source of K+ ions in the 3 M KCl application site; 2) presumed SD-like waves with sequential onset at 2 adjacent recording sites; 3) reversed SD-like waves (rSD) with the spreading sequence between electrodes reversed to the initial one and 4) non-spreading transient depolarizations with ECoG amplitude reduction below 50% on some electrodes, without clear evidence of propagation between them or inconsistent spreading plan (Fig. 8, Supp. Fig. 2). There were no difference between the shape of the amplitude or the time duration between the SDs and PiDs waves.

**Fig. 8.**
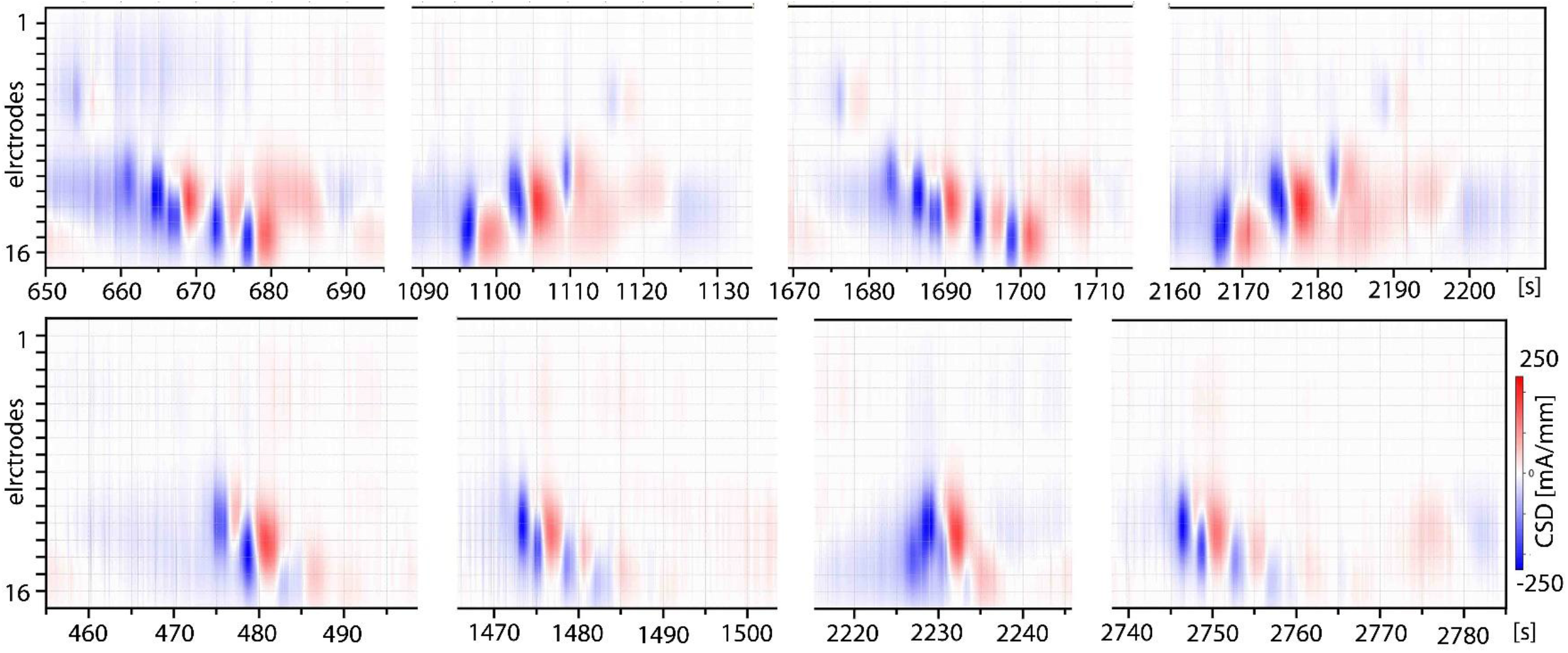
Spread of SDs waves shown with CSD spectrograms. CSD analysis of spreading depolarization events evoked with KCl application. Using kCSD analysis we reconstructed current sources and sinks along electrode positions. We assumed constant tissue conductivity for computational purposes (See in methods - DaSt: Multielectrode recordings). The strongest source was in a location from which the wave came and may have crossed the monitored area in a few different directions (Supp. Fig. 2).

### Protein expression analysis by Western blot (IV)

The ExDP results we obtained were unexpected; we therefore hypothesized that the observed similarities in the protein regulation after SDs and PiDs may be different from those previously suggested by literature data meta-analysis. Since there were no previous experiments testing for differences in protein concentration performed simultaneously on both post-SDs and post-PiDs brain tissues, we analyzed proteins concentration changes in both our experimental models using a comparative western blot paradigm. The chosen proteins, which were shown in previous studiesto influence deprivation-induced cortical representation maps rearrangement, were COX-2 and MMPs (MMP-2, MMP-3, MMP-9; Cybulska-Kłosowicz et al., 2011; Jabłonka et al., 2012; Urbach et al., 2006).

In **C** animals, COX-2 and MMP-3 had a basal level of cortical expression. MMP-2 and MMP-9 in the cortex of **C** animals were barely observed in the somatosensory areas, and in most cases were not present at all. The COX-2 concentration was higher in all experimental groups, but was highest in **SD** somatosensory areas. In the ***BF*** of the **SD** group, the concentration of COX-2 was increased by 5.3 and in **Pt** by 3.2 (Fig. 9). In the ***BF*** of the SD-operated animals, significant increase of all monitored proteins was observed; this was different from the **Pt** group, which showed only COX-2 increase.

**Fig. 9.**
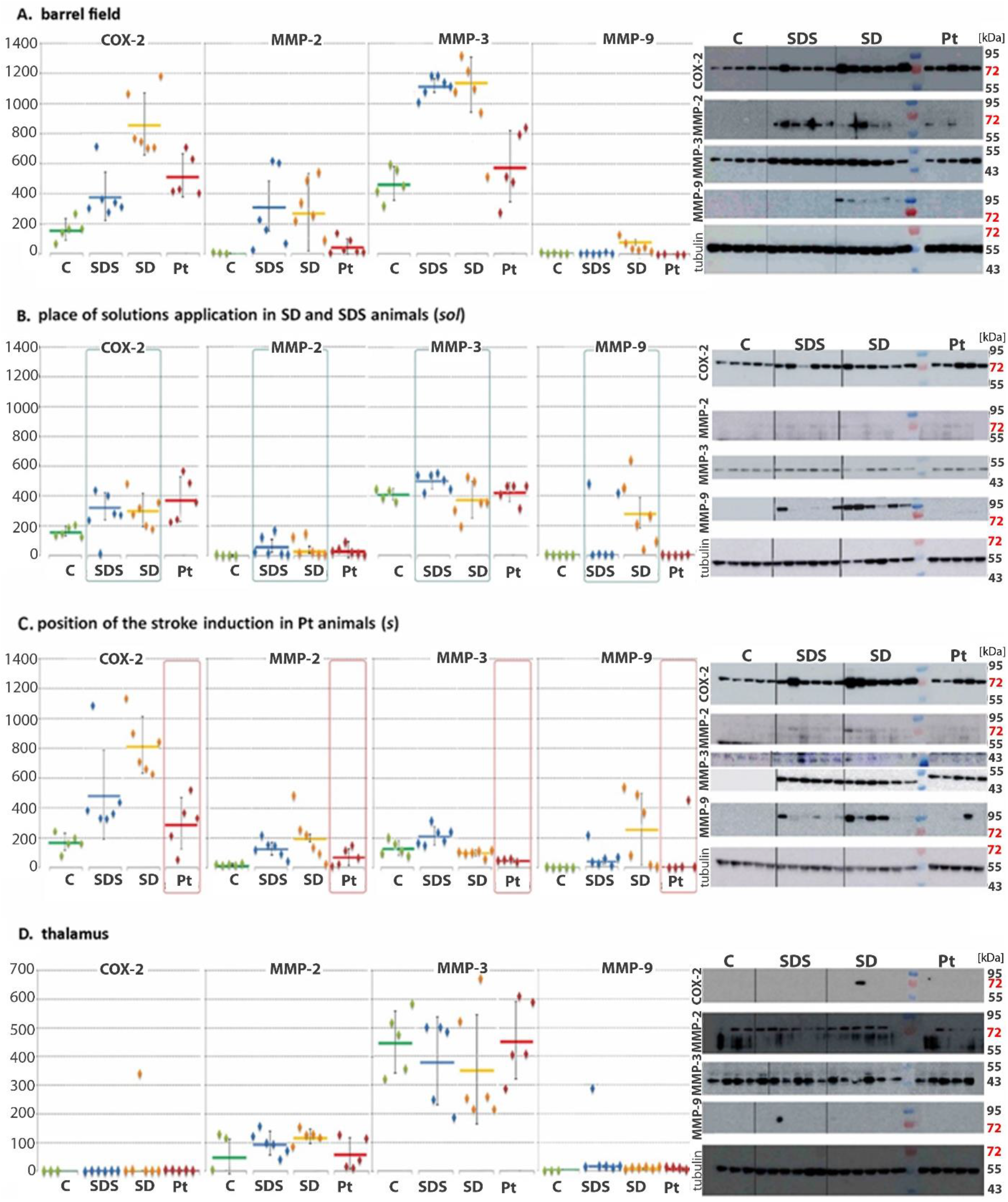
Concentration of COX - 2, MMP - 2, - 3 and – 9 proteins. Western blots of tissue samples from the right hemisphere in experimental animals. Diamonds represent results from individual animals, bar — mean of results for each group. On the right, parts of Western blots photos corresponding to the results shown on the graphs. The lowest blot for each region shows part of the Western blot for tubulin, which was used for normalization. Molecular weight is indicated by molecular weight marker (Supp. Fig. 5). Additional comparisons are shown in Supp. Fig. 6.

In the **SD** group, only COX-2 and MMP-9 increases exceeded the level observed in sham animals. The level of MMP-3 in the ***BF*** increased in all the SD-like operated animals but in other cortical regions it increased only in the **SDS** group. In all monitored cortical areas, MMP-9 increase was most pronounced in **SD** animals.

**C** animals exhibit similar level of the cortex constitutive COX-2 expression in all monitored areas (~150 [An]). The cortical basal interregional comparisons for COX-2 have low deviations (Supp. Fig. 6, Supp Tab1), 11% when expressed as a mean percentage interregional deviation (*PD*). This was different from metalloproteinases, which showed high interregional deviations (MMP-2 and −3: 72%, MMP-9: 107%). Future studies should establish whether the differences are related to some yet undescribed features of MMPs dynamics during post-injury reorganization.

Regarding the thalamic proteins concentration (***th***), COX-2 and MMP-9 were absent in **C** animals but MMP-2 and −3 showed a basal level of expression. MMP-2, −9 and COX-2 appeared in the thalami of all experimental groups. However, the increase was significant only for MMP-2 in the thalami if SDs were performed in the cortex a day earlier. MMP-3 slightly decreased in **SD** and **SDS** animals, but again, not significantly. In almost all the areas the **SD** group seemed to be the most strongly affected.

Additional differences between the groups appeared as different unidentified yet consistent bands recognized by antibodies. We assumed that these unidentified proteins may provide additional information about the mechanisms underlying changes in proteins activities. The cortical modifications of these additional bands were reported for each protein band and are described in the supplementary material.

## Discussion

In this paper we examined the post-SDs and post-stroke cortical plasticity *in vivo* by eliciting partial sensory deprivation of the barrel cortex. Deprivation leads to the enlargement of the cortical representation of the spared whiskers at the expense of the surrounding area (Glazewski and Fox, 1996). Plasticity is impaired in the vicinity of a focal cortical lesion (Jablonka et al. 2007). The pronounced alterations of brain activity following small ischemic injuries affect many aspects of cortical plasticity, including an increase in LTP propensity around the ischemic lesion (Hagemann et al., 1998; Mittmann and Eysel, 2000) and an enhancement in synaptic connectivity that is independent of any purposeful stimulation (Calabresi et al., 2003). Furthermore, the ischemia increases expression of proteins from signaling pathways that have inhibitory roles in axonal outgrowth, like the myelin-associated proteins (Li et al., 2010) or ephrins (Overman et al., 2012).

Our findings confirm that cortical focal stroke lowers ExDP efficiency. PtS in the vicinity BF lowered the widening of the spared whiskers cortical representation surrounded by the deprived cortex (when visualized by 2DG incorporation during bilateral stimulation of spared and homotopic contralateral rows of whiskers). Previous research showed partial restoration of this plasticity impairment when the deprivation onset is delayed after the PtS induction (Jabłonka et al., 2009). The results of this study show that cortical SDs did not influence the expansion of spared whiskers cortical representations after a month of deprivation. Neither the small cortical surface lesion in the occipital cortex nor the SD waves influenced the ExDP in the sensory whiskers representation. This suggests that the impairment in ExDP after a focal cortical lesion is not mediated by SD-like PiDs. What, then, makes these two similar phenomena (SDs and PiDs) so different with respect to metaplasticity changes?

### Alternative mechanisms triggering stroke-induced decrease in cortical plasticity

PtS in the barrel cortex vicinity inhibits plasticity in a COX/PPARs and MMPs dependent manner (Jabłonka et al., 2012; Cybulska-Kłosowicz et al., 2011). The accompanying PiDs are the main factor influencing the post-stroke increase of COX-2 tissue concentration or PPARs activation (Miettinen et al., 1997; Ou et al., 2006; Victor et al., 2006) and subsequent MMPs transcription modifications in the ischemic hemisphere (Candelario-Jalil et al., 2007; Kriz and Lalancette-Hébert 2009). However, we found that the SDs depolarization waves did not change the vulnerability to ExDP cortical map reorganization. We therefore had to consider other factors that could underlie ExDP vulnerability and that would differentiate this two, to some extent similar, experimental groups. We compared electrophysiological and metabolic correlates of cortical activity during and after depolarizations, histological changes, and proteins changes.

### Post injury changes

#### Range of lesions

Both **SD** and **Pt** animals had lesions, but their origin, shape, and range were different. **SD** animals had a surface cortical lesion in the occipital cortex, while **Pt** animals had a columnar lesion in the somatosensory area across all the white matter layers (Jabłonka et al., 2009, 2012). The highly osmotic NaCl solution did not induce depolarizations in SD-operated sham animals (SDS); it produced one-third size cortical surface lesion compared to lesions observed in the **SD** group.

The highly osmotic solutions of NaCl are supposed to cause cellular shrinkage and necrotic cell death, but unlike the case for **SD** animals they do not accelerate additional lesion enlargement (as does treatment with KCl). Therefore, the larger surface lesion size in **SD** vs. **SDS** animals could be treated as a depolarization-induced cell death in the uncomfortable conditions of non-optimal osmotic pressure. Conversely, Pt lesions are induced by regional blood supply restriction, and energetic and metabolic cells failure accompanies PiDs (Song & Yu, 2014). In the experimental model we used the SDs induced lesions in layers II/III of the secondary visual cortex, while the entire cortical columns were destroyed by PtS in a part of the trunk’s sensory representation (Fig 1B, 4B).

The distance between the lesions edge and the B1 barrel was twice as large in **SD** rats as compared to that exhibited by **Pt** animals. PtS experiments demonstrate that the affected Pt lesion core area is more restricted than that shown by other stroke models. Pt is assumed not to induce penumbra effect (Sommer, 2017), spreading the lesion affected area to territories beyond those in which blood flow was reduced. Therefore, since none of the lesions in our experiment should directly affect the ***BF***, we looked for other factors that may influence plastic changes in ExDP after stroke in more distant areas.

### Cortical activity

Spontaneous cortical activity, monitored through electrophysiological recordings and metabolic changes, synchronizes widespread areas and regulates neurites growth. Although PiDs and SDs are generally the same phenomena induced by potassium ions buffering failure (Fabricius et al. 2006, Hartings et al. 2017) the electrophysiological and metabolic modifications and connectivity dysregulation during **Pt** may have different impact on cortical activity homeostasis and/or on interregional wiring (Risher et al. 2010).

SDs require high energy supply, and during SD episode in awake animals, one of largest known cortical 2-DG incorporation occurs (Shinohara et al., 1979). In the post-SDs phases, rapid cerebral blood flow (CBF) increases, followed by a prolonged and sustained CBF decrease (Fabricius et al., 1995).

Our previous work on rats showed long-term bilateral metabolic activity changes after PtS visualized by 2DG incorporation. External stimulation never influenced the metabolic correlates of the cortical activity modified by the injury (which persisted for at least a month) and stabilized two months after stroke (Jabłonka et al., 2009). The inhibitory activity of the hemisphere contralateral to the injured one was documented long ago (Butefisch et al. 2003). Therefore, the increase of the metabolic activity may underlie the interhemispheric inhibition, hypothetically reducing the plasticity changes that follow the stroke. We hypothesized that cortex wiring changed and that metaplasticity was altered under these conditions, in line with Carmichael and Chesselet (2002) findings. These authors found that some experimentally-induced focal cortical lesions can support neurites’ growth and define their direction when the frequency of target area oscillations rhythm was 0.1-0.4Hz. We found no differences between the groups regarding the ECoG signals neither in this frequency nor in other frequencies (Fig. 7 & 8). However, Carmichael and Chesselet’s (2002) observed oscillations within areas functionally connected to the lesion area a few days after the induction of the lesion, while we recorded the signal during the injury episodes, so that may still require further consideration.

We compared the post-stroke and post-SDs metabolic activity by measuring 2-DG incorporation in defined cortical areas one month after the induced episodes. The detected difference in non-stimulated cortical areas disappeared upon comparing the activity relative to the stimulated area of auditory cortex (***Au***), suggesting that just as after Pt stroke, the increase in the spread of 2DG incorporation following SDs was related more to energy-requiring spontaneous activity or physiological metabolic changes than to stimulation-related electrophysiological activity. The interhemispheric differences too disappeared. This means that we observed non-stimulation related interhemispheric changes in metabolic activity a month after the SDs and PiDs episodes. This suggests that the commonly described interhemispheric activity changes that occur after stroke are indeed related to PiDs.

To determine and compare the amplitude and frequency of the depolarizing waves (SDs and PiDs) crossing BF, we used single electrode recordings, which demonstrated spreading waves similarity with higher amplitude of SDs. The multielectrode prefrontal cortex recordings showed radial flow from the core locus of the treatment after applying KCl solution onto the occipital area. However, reversed and irregular waves without a defined trajectory were also observed, in line with recent evidence (Kaufmann et al., 2016) that showed less SDs homogeneity than previously described. That may require a further validation using ECoG recordings at the place of cortical plasticity origin. Especially, that not only the SDs propagation variety was described within the cortex but also some features differing cortical regions propensity to SD generation and propagation that indicate a special vulnerability of BF to SD generation when the extracellular potassium concentration increases (Bogdanov et al. 2016).

### Interhemispheric equilibrium

SDs are usually unilateral symptoms (Goadsby 2001). In this study we show they also affect the contralateral, “intact” hemisphere (Fig. 5). It seems obvious that any activity disturbances have to influence contralateral network functioning since the activity equilibrium is altered (Vecchia & Pietrobon, 2012); a unilateral injury is indeed considered as a source of entire brain activity imbalance (Mun-Bryce et al. 2004). However, can a unilateral injury modify entire brain metaplasticity, thus changing the dynamics of cortical map remodeling? Recent reports suggested that unilateral plasticity changes may bilaterally modify SII representations (Debowska et al. 2011), in a similar way to the effect reported in this paper. On the other hand, the diaschisis in the injured hemisphere leads to prolonged increase of metabolic markers of cortical activity (like 2DG or fMRI images) in the “intact” hemisphere (Dijkhuizen et al. 2003, Jablonka et al. 2010). This increase is not related to activity induced by sensory stimulation (Jablonka et al. 2010). Nevertheless, no studies, to date, have investigated demonstrated directly the source of the increase in cortical activity and its influence on metaplasticity.

It was suggested that spontaneous SDs and migraine might be an effect of inhibitory and excitatory imbalance (Vecchia & Pietrobon, 2012). If this is the case, both should influence ExDP. In our studies, however, the repeated SDs, although generating the prolonged activity increase in the “intact” hemisphere, did not modify cortical map rearrangement triggered by the ExDP. On the other hand, the imbalance between the hemispheres reflected in larger 2DG incorporation in the intact hemisphere than in the treated one was larger after the stroke and was the only measured parameter that pronouncedly differentiated distingsuishedthe **Pt** and the **SD** groups. Both in the poststroke and post SDs animals, the imbalance was independent of sensory stimulation, suggesting that similar but stronger mechanisms are operating after focal cortical ischemia than after SDs. This might be due to deleting part of the neuronal net module in the stroke core, the absence of which may modify areas previously connected to the destroyed region. The homotopic contralateral areas undoubtedly belong to this category. The prolonged increase in the activity in the “intact” homotopic region might be differently balanced after stroke, influencing the BF via callosal or subcortical connections (Mun-Bryce et al. 2004). Further studies are required to support this suggestion.

### Proteins

Data obtained by Urbach et al. (2006) showed 1.6-fold increase in COX-2 mRNA in the somatosensory cortex 24h after SDs induction by KCl. This increased expression persisted until the 7th day. Other studies also detected increased COX-2 protein up-regulation within 24h after SDs episodes (Bidmon et al., 2000; Koistinaho et al., 1999; Thomson and Hakim, 2005). Hence, there are similarities in COX-2 increase following Pt focal lesion and SDs (Yin et al. 2002, Urbach et al. 2006).

Why, then, does SD inflammatory induction fail to inhibit plasticity (Jander et al., 2001), while the post-stroke COX-2/PPARs and/or MMPs induction does inhibit it (Cybulska-Kłosowicz, 2011 Jabłonka et al., 2012)? SDs/PiDs were assumed to be mainly responsible for transcriptional changes in areas “remote from” the cortical injury locus (Hermann et al., 1999; Küry et al., 2007; Miettinen et al., 1997). In our experiments the distance of the stroke-induced lesion edge from the closest B1 barrel edge was 1.2 mm. The furthest barrel of the non-deprived row B was almost 3 mm away, which can certainly be described as the “remote non-ischemic cortex” in which cortical spreading depressions were identified as the main mechanism underlying the protein expression changes (Küry et al., 2004).

The PiDs-related changes of the protein expression profile after focal stroke in the remote areas of the affected hemisphere and the changes accompanying SDs are very similar (Cybulska-Kłosowicz, 2011; Gursoy-Ozdemir et al., 2004; Urbach et al., 2006). Although some of the studies that explored spreading depolarizations investigated the effects of ischemic injury (Anderson et al., 2003; Chapuisat et al., 2008; Dreier, 2011), no previous experiments compared SDs and PiDs in the same experimental set up with proteins *compared on the same blots*. There were no previous comparisons of the spatial expression profiles in SDs and PiDs. We compared protein levels at several cortical regions that were either closer or more distant from the treated areas as well as thalamus.

#### Base level of COX-2, MMP-2, −3, −9 in monitored areas

Contrary to other tissues, the brain’s COX-2 is constitutively expressed, but both constitutive and inducible pools participate in synaptic plasticity and facilitate electrophysiological and behavioral symptoms of cognitive functions (Sang and Chen, 2006). The matrix metalloproteinases, including MMP-2, MMP-3, and MMP-9, are synthesized and secreted as inactive proenzymes (zymogen), which are enzyme-activated in the extracellular space through different MMPs domain interactions (Djuric and Zivkovic, 2017; Woessner, 1991).

The cortical synthesis of MMP-3 and COX-2 in the **C** group (Fig. 9 & 10; Supp. Fig. 5), confirm the constitutive expression of these two proteins in rats that was described for human brains (Kirkby et al., 2016), and both participate in the establishment of cortical plasticity. Yet, MMP-2 and MMP-9 were not detected in any of the selected **C** areas. However, since MMP-9 was expressed at a low level in healthy brain synapses (Huntley, 2012; Reinhard et al., 2015), it was probably below our Western Blot detection threshold (as previously shown in mice; see Wang et al., 2000). In the thalamus (***th***) of mice, COX-2 mRNA and MMP-9 protein and its activity were undetected (Samad et al., 2001; Wang et al., 2000). This was confirmed in this study. However, we have shown constitutive expression of the active form of MMP-2 (64kDa) and MMP-3 (53kDa) in the ***th*** of the **C** group.

#### Post-stroke and post-SDs, protein changes

Although COX-2, MMPs, and PPARs influence the lesion size after middle cerebral artery occlusion (Gu et al., 2005), no changes in the infarct core volume were observed after ibuprofen or FN-439 treatment in our focal photothrombotic stroke model (Cybulska-Kłosowicz, 2011; Jabłonka et al., 2009, 2012). Therefore, it is improbable that COX-2 or MMPs or PPAR-gamma inhibition facilitate the lesion-induced necrosis size. It is probably due to the lack of penumbra in PtS stroke model (Sommer, 2017). This suggests that direct cells necrosis does not influence plasticity after stroke.

COX-2 was the only protein in the ***BF*** whose concentration was increased following PtS with PiDs. Our results show that MMP-9 is not the most strongly activated metalloproteinase after focal cortical stroke. In ***BF***, it was detected only after SDs and had much lower expression than MMP-2 and −3, which were both up-regulated by high molarity solution applications.

MMP-2 zymogen (72kDa) was present mainly in the treated animals and the bands were most prominent in ***s*** of **SDS** and **SD** animals. In ***BF***, the MMP-2 bands were only present in the active form of 64kDa. **SD** and **SDS** groups had the strongest bands of the active form of MMP-2. Interestingly, MMP-2 was most pronounced in ***BF***, although the regions closer to the treated area had a lower level of the active form of MMP-2 and even less than the zymogen, which was three times lower than in the ***BF*.** The most pronounced MMP-2 activation was seen in the ***BF*** of **SDS** animals. Thus, the solution application and the subsequent ions equilibrium changes may have triggered MMP-2 zymogen synthesis and high-level activation in the ***BF*** but not necessarily in the surrounding cortex, which might be less vulnerable to extracellular ion changes (Bogdanov et al. 2016). Thalamic changes in all groups were less pronounced (Supp.2, and Supp.4).

Comparisons of the amount of protein after the SDs and focal stroke with PiDs showed no relation between COX-2, MMP-2, −3 (Supp. 5) and −9 and ExDP. This suggests that higher protein levels in the ***BF*** are not the factor inhibiting the widening (as visualized by 2DG), of the spared whiskers representation after stroke, as all the changes were at least doubled in the **SD** group, which had no influence on ExDP. However, even a slight upregulation of MMP-2 or −9 in the synapses of the BF could protect synaptic plasticity (Gontier et al. 2009, Michaluk et al. 2011) and therefore maintain the level of ExDP after the SDs (Kaliszewska et al. 2011).

The higher metalloproteinases concentration in the **SD** group (compared to the **Pt** group) could explain the observed changes in metaplasticity. This suggests an existence of competitive mechanisms which balance and compensate the plasticity-supporting changes in post-injury tissue rearrangement.

### Concluding remarks

Our study shows that changes in metalloproteinases (MMP-2, −3, −9) and COX-2 levels are not responsible for experience-dependent plasticity impairment after focal cortical stroke. The global changes in brain functioning accompanying the spreading depressions are similar to those observed after focal Pt lesion with periinfarct depolarizations. Plasticity in the ***BF*** however, was diminished only after focal cortical stroke. The SDs have no negative influence on cortical experience dependent plasticity so neither the PiDs nor the inflammatory response triggering the protein changes are the main causes of cortical metaplasticity changes after focal cortical stroke.

COX-2 was the only protein which increased in BF one day after stroke in its vicinity. Yet, specific to **SD** MMP-9 concentration increase could have a protective influence stabilizing the metaplasticity. Nonetheless, the COX-2 and MMP-9 increase in BF were still most pronounced following SDs. Slight increase of MMP-9 in the thalamus was observed in all the animals with focal injury in the cortex (no matter the surface or entire cortical column) which might suggest some kind of thalamic reorganization.

The PiDs amplitude of the depolarization was a bit lower than SDs but no other differences were observed in duration and frequencies of spontaneous oscillations during the depression period. The spreading waves of SDs had no radial distribution within the cortex which suggest it spreads diversely within the cortical areas.

Our study shows that spreading depressions have a global influence on the brain functioning and induce not only the protein expression changes in entire depolarized hemisphere but also in thalamus. Pronounced dysregulation of cortical activity and interhemispheric activity equilibrium was observed even a month after repetitive spreading depression episodes

Although the changes observed in our experimental models are usually more pronounced after SDs, the interhemispheric metabolic activity ratio was an exception (Fig. 5), and was higher after stroke. Hence, the interhemispheric interactions modified by stroke may be a promising target for future studies of post-stroke experience-dependent plasticity and convalescence.

## Availability of data and materials

The manuscript data will be stored at the Research Gate profile of the corresponding author.

## Acknowledgements

We would like to thank Prof. Marta Wiśniewska for her help in protein studies and Prof. Ewa Kublik for her help in electrophysiology and especially Dr. rer. nat. Anja Urbach and Prof. Otto Witte for their continuous support and sharing their data and to Prof. Małgorzata Kossut for supporting the scientific development. We would also like to thank Dr Elżbieta Fuszara and Magdalena Grabowska for a lot of help with setting up the experimental paradigm, equipment and data analysis.

Jan Jabłonka and collaborators were funded by grants: MNiSW 1008/PO1/2007/22; 0184/IP1/2011/71, 0292/IP1/2013/72.

Paulina Urban was funded by grant number NCN 2014/15/B/ST6/05082.

## Author Contributions

Maria Sadowska has done protein analysis under the supervision of Lukasz Szewczyk and and contributed to the writing of the manuscript. Clemens Mehlhorn helped in setting up 2DG brain mapping and the statistical analysis of the cortical whiskers representation. Władysław Średniawa and Paulina Urban did the electrophysiological data analysis and part of the whiskers cortical representation analysis. Maciej Winiarski has done the interhemispheric comparisons of the 2DG incorporation. Aleksandra Szlachcic helped at all the stages of the study. All the experimental stages were supervised by Jan Jablonka, who wrote the manuscript.

## Supplementary materials

### Experience-dependent plasticity (I)

#### Supp.1

One month of unilateral deprivation of all but one row of whiskers resulted in non-deprived rows widening in all experimental and control groups [Fig. 2 & 3]:

- **C:** in the control group, deprivation resulted in 42% ± 14% widening of non-deprived row B cortical sensory representation in the hemisphere contralateral to the deprived whiskers (1160 ± 121 μm) in comparison to the same row in the non-deprived hemisphere (814 ± 15 μm; p = 0.009);
- **Pt:** after stroke the deprivation resulted in 11% ± 9% widening of the non-deprived row B cortical sensory representation (906 ± 76 μm to 821 ± 113 μm; p = 0.017);
- **SDC:** in aCSF treated group deprivation resulted in 35% ± 27% non-deprived row cortical sensory representation widening (1075 ± 132 μm) in comparison to the same row in the non-deprived hemisphere (798 ± 108 μm; p = 0.03);
- **SDS:** in the sham group, 3M NaCl application resulted in a small lesion in occipital cortex. The difference in the width between rows B representations of the deprived (1023 ± 192 μm) and non-deprived (779 ± 123 μm) hemispheres was 33% ± 26%, p = 0.04;
- **SD:** in animals with spreading depolarisations and focal lesion induced by 3M KCl the difference between rows B representations was significant (p <0.001). However, the mean percentage difference between deprived and non-deprived sites was 42 % ± 18% like in intact control group C; (1036 ± 192 μm in deprived hemisphere vs. 781 ± 81 μm in non-deprived; p = 0.002).

The stroke group was the only one that differed significantly from others when the widening was compared (interhemispheric difference of the rows B representations width 11% to: 42% in **C**, p=0.002; 35% in **SDC**, p=0.009; 33% in **SDS**, p=0.004 and 42% in **SD**, p=0,006). The non-deprived row B representation in the deprived hemisphere with focal PtS in the **Pt** animals differed significantly from the homologus areas in other groups: **SD** (p=0.006), **C** (p=0.003) and **SDC** (p=0.04). There were no statistically important differences between SD-operated groups in the non-deprived row B cortical representations [Fig. 3].

### Proteins (I)

#### Additional epitopes

COX-2 bands 72 kDa were not observed in the thalamus. However, an epitope on bands clustered around 180kDa was detected in **C**, **Pt** and two **SDS** (Supp.2). MMP-2 was not detected in the intact **C** animals neither in the thalamus nor in the cortex. (Supp.3).

In the thalamus, as in the cortex, the full-length MMP-3 had 45 kDa bands. MMP-9 in the **C** animals were undetected in the thalamus and the cortex. In the thalamus MMP-9 was only present in one **SDS** animal (Supp. 4) However, an additional epitope of 26kDa bands of MMP-9 was present in the thalamus of all experimental groups but not in the control groups. The epitope was detected just in the thalamic probes, suggesting a different damage mechanism from that observed in the cortex. Thus, it is not only stroke that induces thalamic changes through retrograde signaling or even cell death. In our experimental models both SD-operated groups modified the thalamic functioning involving the metalloproteinases and COX-2 changes. That could suggest that even surface lesions induce subcortical modifications reflected in MMP-9 epitope representation in the thalamus.

Interestingly, two of the three **SD** animals which had MMP-2 in ***th*** had also increased active form of MMP-2 (64kDa) concentration in the cortex’s ***BF*** region. This suggests that the MMP-2 activity is affected by the changes of the afferent regions. It is in line with results showing MMP-2 increase, extracellular transfer and colocalization with GAP-43 in an area with anterograde connections from injured cochlea (Fredrich and Illing 2009). Gontier et al (2009) suggested that MMP-2 enables synaptogenesis to the GAP-43 but it may also release neurites from extracellular semaforins and reinforce the dendritic trees rearrangement. Therefore, MMP-2 “the major actor in extracellular matrix recomposition” should support the cortical map remodeling in ExDP.

#### Supp. 2

Thalamic changes in all groups were less pronounced, and COX-2 (bands 72 kDa) were not observed. However, an epitope of bands clustered around 180kDa was detected with the highest concentration in four out of six animals of the **SD** group (86±24). Two animals from **Pt** group demonstrated the equivalent level of these bands in the ***th*** (61±19), but other animals did not. Conversely, the **SDS** animals had a relatively low deviation (PD=45%), although the level of the bands OD had a much lower threshold (21±14). We assumed that it was a procedural artifact; nonetheless, the **SD** had a significantly higher density of COX-2 (180kDa, 63±40 vs. **C** 6±6, p=0.02 vs. sham 21±13, p=0.04) and shams had a significantly higher concentration in that band than **C** (p=0.04).

#### Supp. 3

One of **C** animals had MMP-2 zymogen detected and two had the unidentified 45kDa epitope which was present in the other experimental groups. The 45kDa epitope was shown to be specific for different anti-MMP-2 antibodies (Novus Biotechnology MMP-2; 43 kDa, cat number - NBP2-37426). Significantly higher concentration of the 45kDa epitope was detected in three animals after stroke (323±24 [An] while in all the other animals in all the experimental groups the concentration was 198±56, p=0.0009).

#### Supp. 4

MMP-9 in C animals was not detected neither in the thalamus nor in the cortex. MMP-9 in the thalamus was present just in one **SDS** animal, the same animal that had higher MMP-2 thalamic level (157 [An] vs. 97±40 [An] in the rest of the SDS) and was one of the two animals that had lower MMP-3 (248 [An] vs. 496±9 [An] for the rest three of SDS).

### Supplementary Figures

**Supp. Fig. 1:**
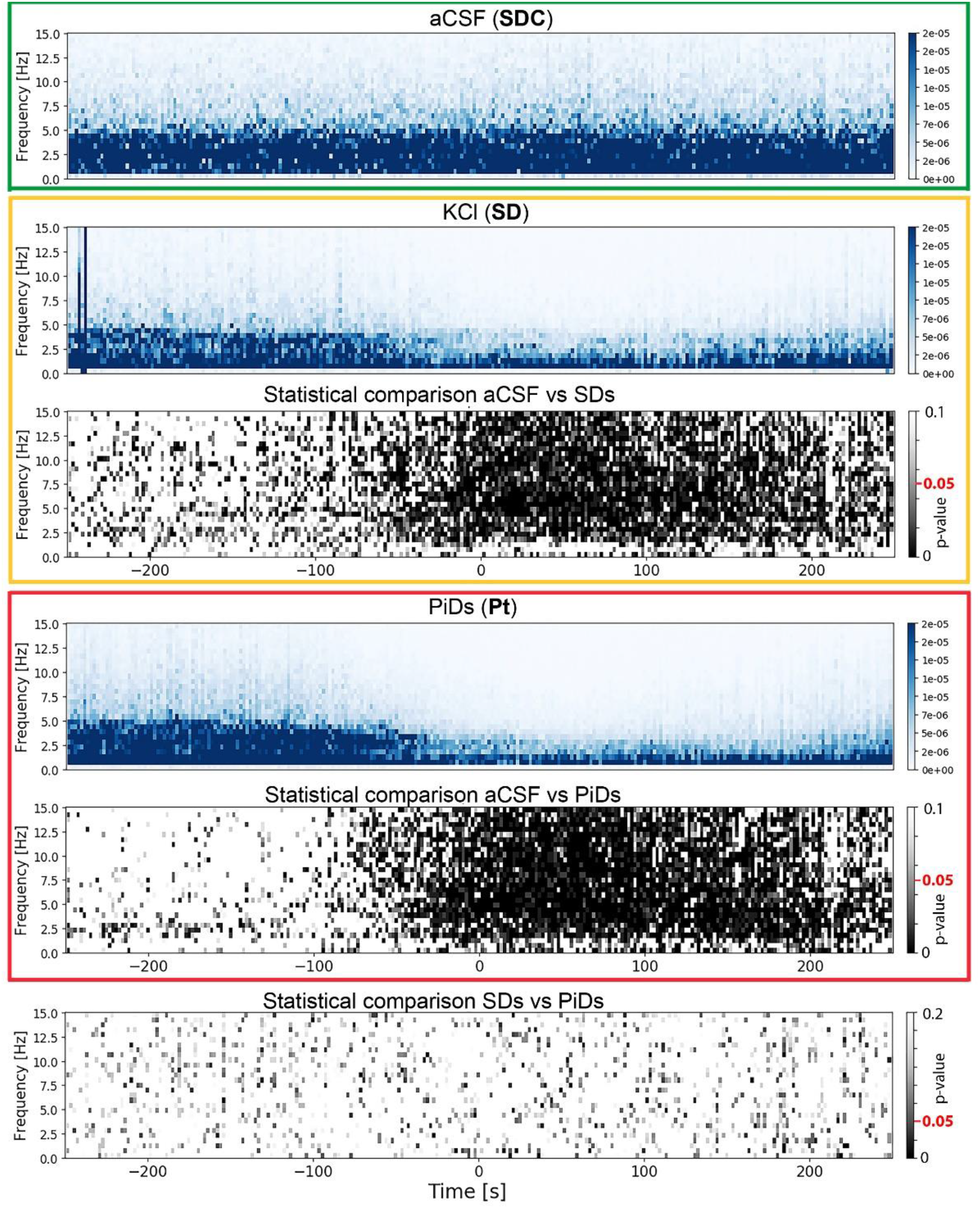
**The ECoGs spectrogram** matrices (blue) and statistical comparison (black) between baseline, aCSF treated, **SDC** animals (green frame), SDs of **SD** animals (orange frame) and PiDs of **Pt** (red frame). Timeline is related to through of single SD event (timepoint 0 – minimal voltage value; ±200 [s]). The estimation of difference significance between groups was calculated using a resampling method. (See Methods - DaSt: Electrophysiological statistic). Every square represent p-value for a difference between the means for new random sets of fragments and the original difference between two groups (the gray scale of the spectrograms). During the course of a depolarization wave there was a significant difference between the baseline of **SD** and **Pt** versus **SDC** (p=0.03) There was a significant reduction of 2-12 Hz oscillations during depolarization both for SDs and PiDs but PiDs and SDs did not differ from each other. Spreading depolarizations probably causes sedation of the large cell populations.

**Supp. Fig. 2:**
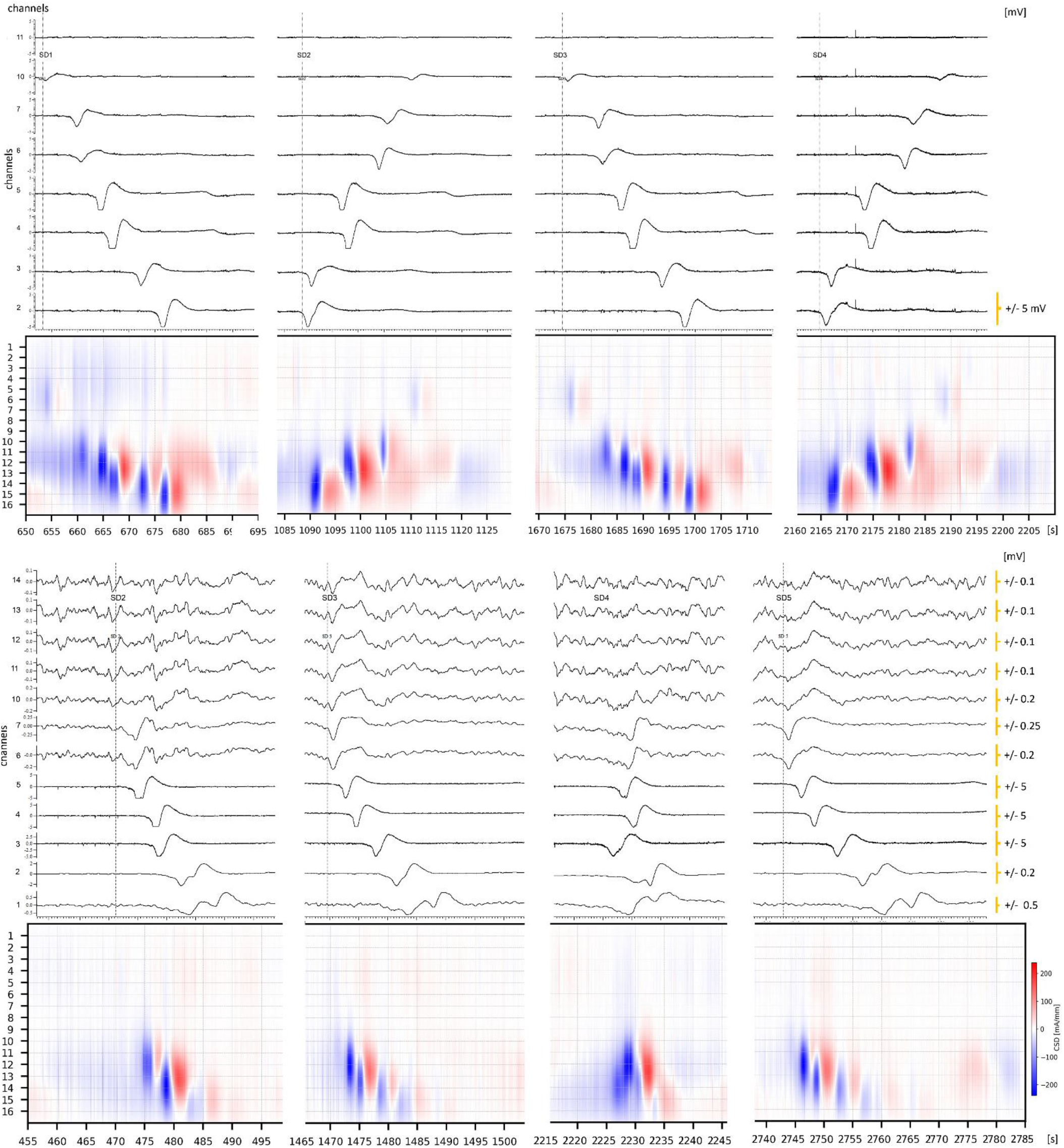
Spreading waves and CSD for spreading waves of SDs. LFP and CSD analysis of spreading depolarization events evoked with KCl application in two different animals. Using kCSD analysis we reconstructed current sources and sinks along electrode positions (See in methods - DaSt: Electrophysiological statistic). We assumed constant tissue conductivity for computational purposes. The strongest source was in the location from which the wave originated. As can be seen it crossed the monitored area in several directions for the same animal.

**Supp. Fig. 3:**
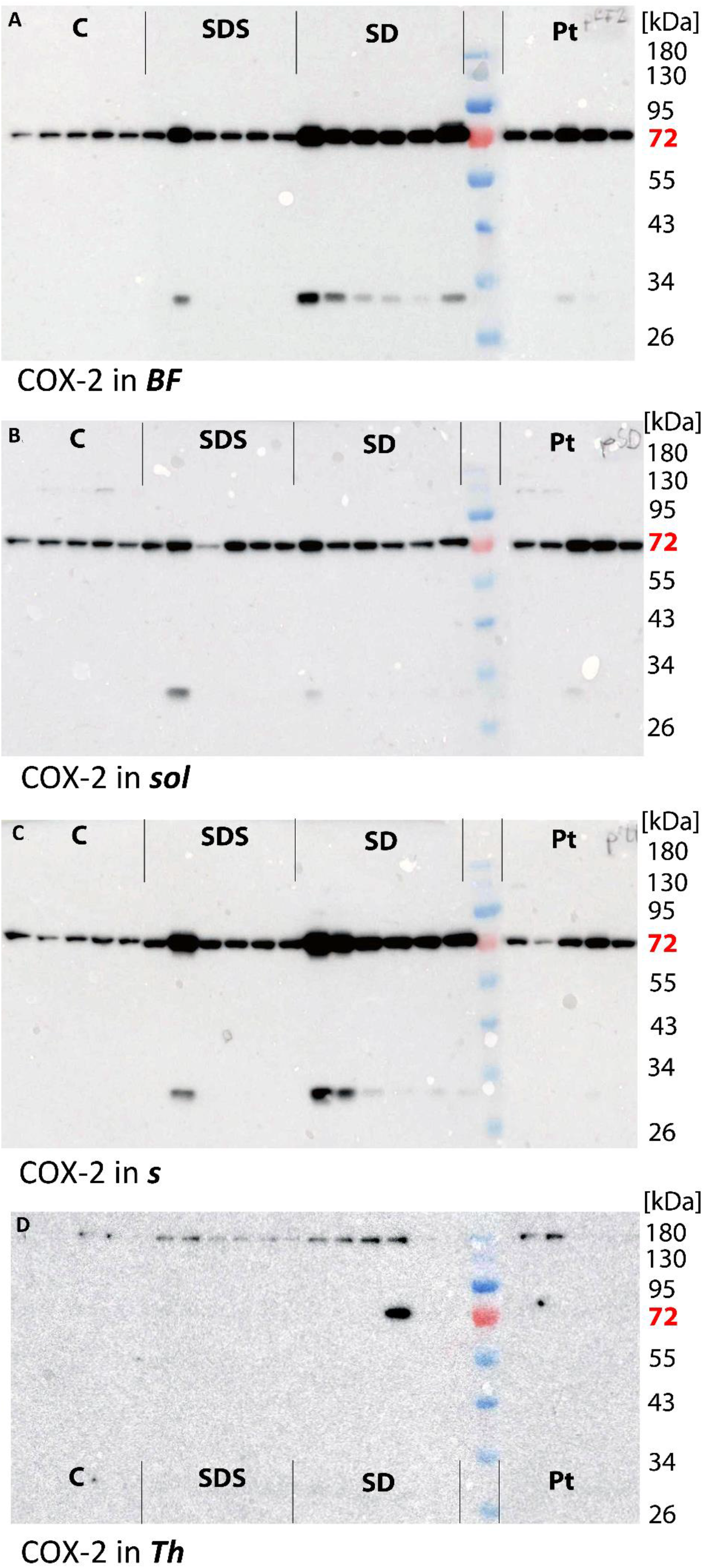
Western blots from brain tissue samples incubated against COX-2. In the cortical areas of (A) barrel fields, (B) the place of solution application, (C) the position of the stroke induction and (D) thalamus with visible additional bands. The protein degradation product (33kDa) is related to the higher concentration of COX-2 in one specimen. In the thalamus series of 180 kDa bands are the only presented for the whole animal series but increased in experimental groups.

**Supp. Fig. 4:**
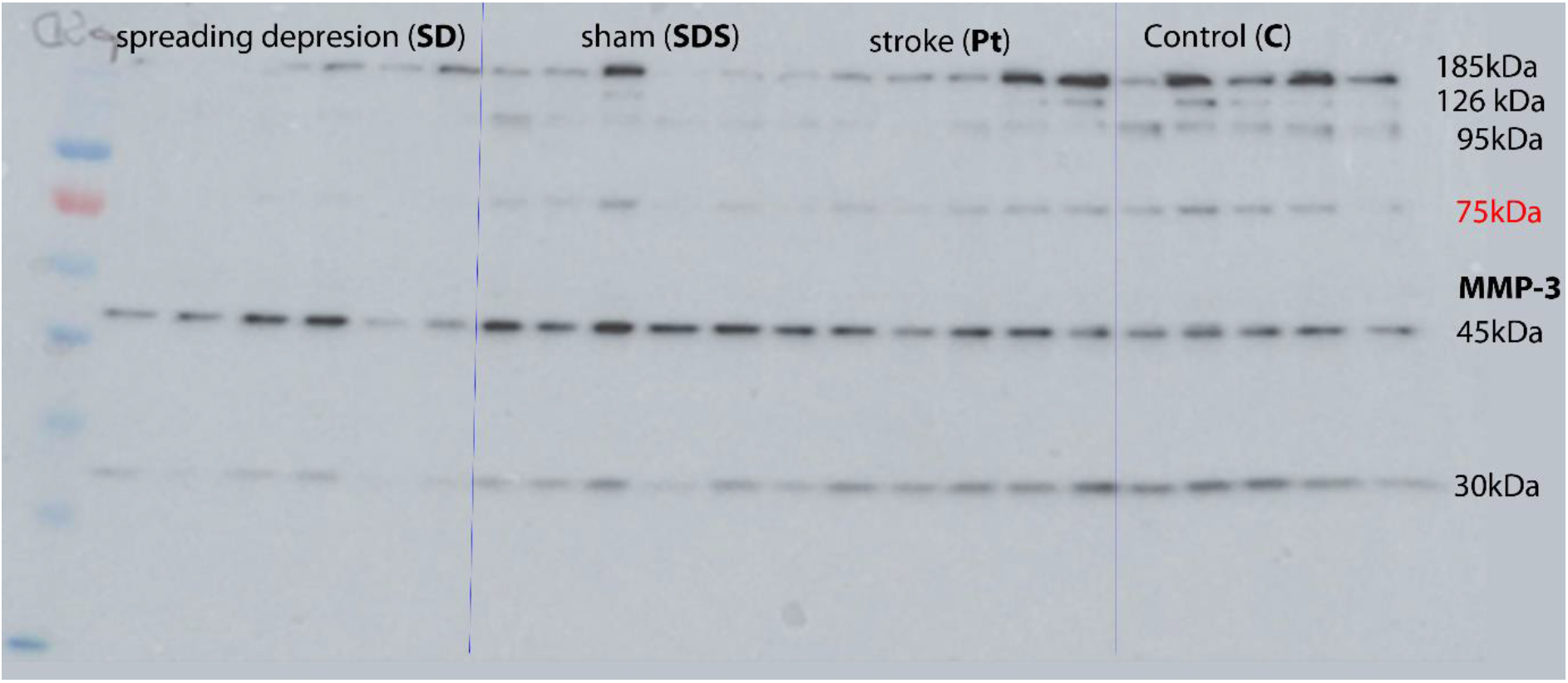
Western blot membranes with protein lysates from brain tissue samples incubated against MMP-3. Additional bands are visible in SD-operated animals, in the cortical area where the solution was applied. The protein degradation product (30kDa) is due to the lower concentration of MMP-3 in the group of **SD** animals.

**Supp. Fig. 5:**
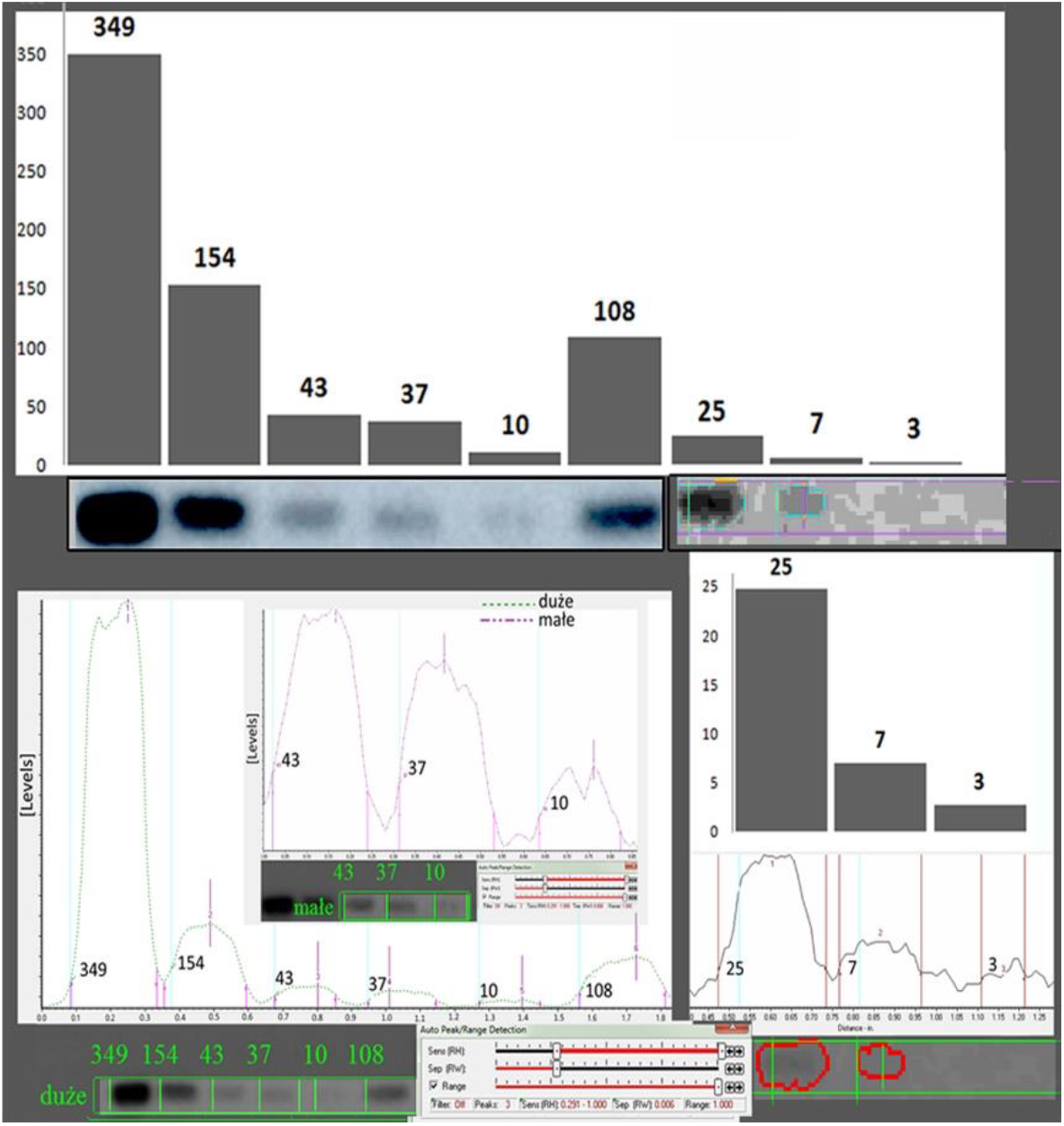
Western blot analysis. Top: illustration of the data acquisition procedure, the reference bands of protein different concentration and the relative abstract number [An] (respective values of OD multiplied by band surface and divided by reference housing protein multiplied by 1000). Bottom: line analysis derived from the MCID program designed for electrophoresis with “peak detection” and “smiling correction”. The series of bands on a bottom with OD and measurement of bands areas.

**Supp. Fig. 6.**
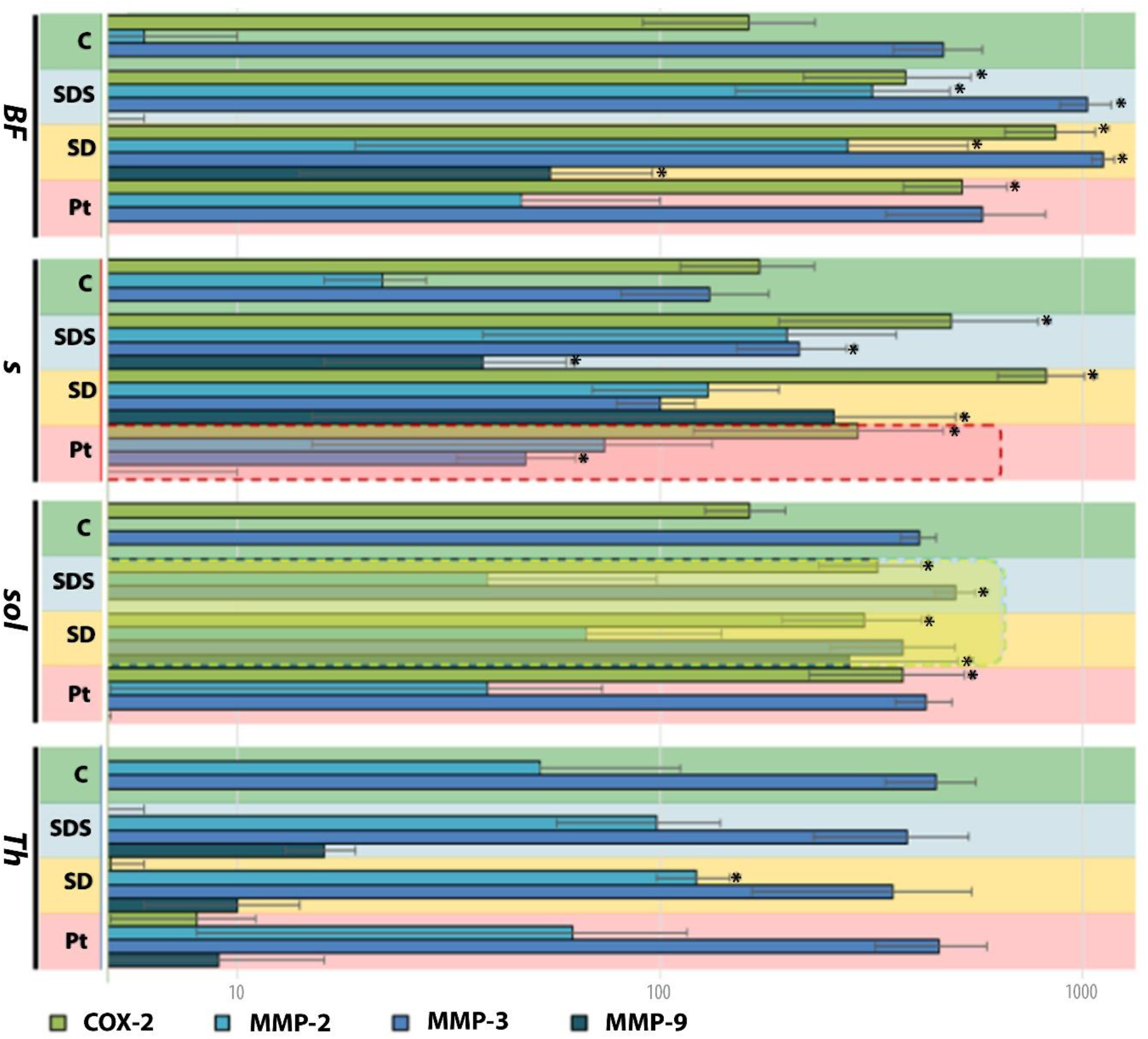
Concentration of COX-2, MMP-2, −3, −9 by Western blots for all the groups in different regions. A logarithmic scale of the abstract number [An] derived from OD comparison. The asterix point to significant difference from the control level of the protein concentrations (p<0.05).

**Supp. Tab. 1:**
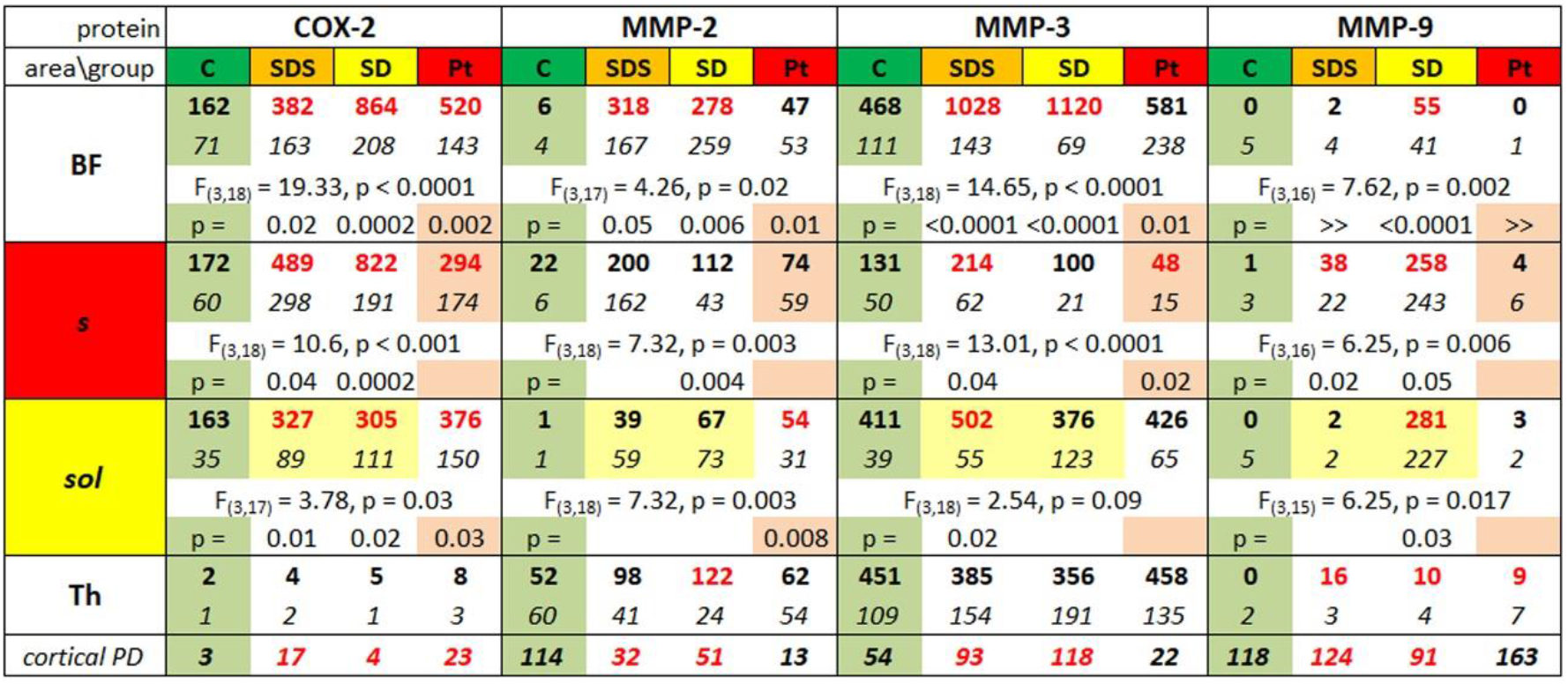
Protein concentration in abstract numbers [An] (An of a band OD minus the OD of the reference housekeeping protein [levels] x area [um]) with *SD* below. Values significantly different from control are in red; p < 0.05. Percentage deviations of cortical proteins amount on the bottom part show that metalloproteinases had ten times higher deviation than COX-2 in the intact brains and from 3 to 7 times higher postoperative intracortical deviations. Green background marks the control values. The injured areas are marked with color (yellow – the highly osmotic solution application, red – the illuminated area injured by the PtS).

## References

Anderson MF, Blomstrand F, Blomstrand C, Eriksson PS, Nilsson M. Astrocytes and stroke: networking for survival? Neurochemical research. 2003 Feb 1; 28(2):293–305.

Andrews RJ. Transhemispheric diaschisis. A review and comment. Stroke. 1991 Jul; 22(7):943–9.

Butefisch CM, Netz J, Wessling M, Seitz RJ, Homberg V. Remote changes in cortical excitability after stroke. Brain 2003; Mar; (3)126:470–81.

Bidmon HJ, Oermann E, Schiene K, Schmitt M, Kato K, Asayama K, Witte OW, Zilles K. Unilateral upregulation of cyclooxygenase-2 following cerebral, cortical photothrombosis in the rat: suppression by MK-801 and co-distribution with enzymes involved in the oxidative stress cascade. Journal of chemical neuroanatomy. 2000 Nov 1; 20(2):163–76.

Bogdanov VB, Middleton NA, Theriot JJ, Parker PD, Abdullah OM, Ju YS, Hartings JA, Brennan KC. Susceptibility of primary sensory cortex to spreading depolarizations. Journal of Neuroscience. 2016 Apr 27; 36(17):4733–43.

Calabresi P, Centonze D, Pisani A, Cupini LM, Bernardi G. Synaptic plasticity in the ischaemic brain. The Lancet Neurology. 2003 Oct 1;2(10):622–9.

Candelario-Jalil E, González-Falcón A, García-Cabrera M, León OS, Fiebich BL. Post-ischaemic treatment with the cyclooxygenase-2 inhibitor nimesulide reduces blood-brain barrier disruption and leukocyte infiltration following transient focal cerebral ischaemia in rats. Journal of neurochemistry. 2007 Feb;100(4):1108–20.

Carmichael ST, Chesselet MF. Synchronous neuronal activity is a signal for axonal sprouting after cortical lesions in the adult. Journal of Neuroscience. 2002 Jul 15;22(14):6062–70.

Chapuisat G, Dronne MA, Grenier E, Hommel M, Gilquin H, Boissel JP. A global phenomenological model of ischemic stroke with stress on spreading depressions. Progress in biophysics and molecular biology. 2008 May 1;97(1):4–27.

Charles AC, Baca SM. Cortical spreading depression and migraine. Nature Reviews Neurology. 2013 Nov;9(11):637.

Cybulska-Kłosowicz A, Liguz-Lecznar M, Nowicka D, Ziemka-Nalecz M, Kossut M, Skangiel-Kramska J. Matrix metalloproteinase inhibition counteracts impairment of cortical experience-dependent plasticity after photothrombotic stroke. European Journal of Neuroscience. 2011 Jun;33(12):2238–46.

Debowska, W., Liguz-lecznar, M., & Kossut, M.. Bilateral Plasticity of Vibrissae SII Representation Induced. Journal of Neuroscience. 2011 Apr. 5989-10.2011

Dienel GA, Hertz L. Astrocytic contributions to bioenergetics of cerebral ischemia. Glia. 2005 Jun;50(4):362–88.

Dietrich WD, Ginsberg MD, Busto R, Smith DW. Metabolic alterations in rat somatosensory cortex following unilateral vibrissal removal. Journal of Neuroscience. 1985 Apr 1;5(4):874–80.

Dihné M, Grommes C, Lutzenburg M, Witte OW, Block F. Different mechanisms of secondary neuronal damage in thalamic nuclei after focal cerebral ischemia in rats. Stroke. 2002 Dec 1;33(12):3006–11.

Dijkhuizen RM., Singhal AB, Mandeville JB, Wu O, Halpern EF, Finklestein SP, Lo EH. Correlation between brain reorganization, ischemic damage, and neurologic status after transient focal cerebral ischemia in rats: a functional magnetic resonance imaging study. The Journal of Neuroscience: The Official Journal of the Society for Neuroscience, 2003 May 23(2), 510–517.

Djuric T, Zivkovic M. Overview of MMP Biology and Gene Associations in Human Diseases. The Role of Matrix Metalloproteinase in Human Body Pathologies. 2017 Dec 20:1.

Dreier JP. The role of spreading depression, spreading depolarization and spreading ischemia in neurological disease. Nature medicine. 2011 Apr;17(4):439.

Drew PJ, Feldman DE. Intrinsic signal imaging of deprivation-induced contraction of whisker representations in rat somatosensory cortex. Cerebral Cortex. 2008 May 30;19(2):331–48.

Ethell IM, Ethell DW. Matrix metalloproteinases in brain development and remodeling: synaptic functions and targets. Journal of neuroscience research. 2007 Oct;85(13):2813–23.

Fabricius MA, Akgoren NU, Lauritzen M. Arginine-nitric oxide pathway and cerebrovascular regulation in cortical spreading depression. American Journal of Physiology-Heart and Circulatory Physiology. 1995 Jul 1;269(1):H23–9.

Fabricius, M., Fuhr, S., Bhatia, R., Boutelle, M., Hashemi, P., Strong, A. J., & Lauritzen, M. Cortical spreading depression and peri-infarct depolarization in acutely injured human cerebral cortex. Brain. 2006 Aug 129, 778–790.

Faraguna U, Nelson A, Vyazovskiy VV, Cirelli C, Tononi G. Unilateral cortical spreading depression affects sleep need and induces molecular and electrophysiological signs of synaptic potentiation in vivo. Cerebral Cortex. 2010 Mar 26;20(12):2939–47.

Footitt DR, Newberry NR. Cortical spreading depression induces an LTP-like effect in rat neocortex in vitro. Brain research. 1998 Jan 19;781(1-2):339–42.

Fredrich M and Illing RB. MMP-2 is involved in synaptic remodeling after cochlear lesion. Regeneration and transplantation. NeuroReport. 2010 Nov 21:324–327

Fujioka H, Dairyo Y, Yasunaga KI, Emoto K. Neural functions of matrix metalloproteinases: plasticity, neurogenesis, and disease. Biochemistry research international. 2012;2012.

Fukuda S, Fini CA, Mabuchi T, Koziol JA, Eggleston Jr LL, del Zoppo GJ. Focal cerebral ischemia induces active proteases that degrade microvascular matrix. Stroke. 2004 Apr 1;35(4):998–1004.

Glazewski ST, Fox KE. Time course of experience-dependent synaptic potentiation and depression in barrel cortex of adolescent rats. Journal of neurophysiology. 1996 Apr 1;75(4):1714–29.

Glazewski S, Giese KP, Silva A, Fox K. The role of α-CaMKII autophosphorylation in neocortical experience-dependent plasticity. Nature neuroscience. 2000 Sep;3(9):911–8.

Goadsby PJ. Migraine, aura, and cortical spreading depression: why are we still talking about it?, Ann. Neurol. Jan 49 (2001) 4–6.

Gonthier B, Koncina E, Satkauskas S, Perraut M, Roussel G, Aunis d, Kapfhammer JP0, Bagnard D. A PKC-Dependent Recruitment of MMP-2 Controls Semaphorin-3A Growth-Promoting Effect in Cortical Dendrites. 2009 PLoS One. 2009; 4(4): e5099.

A PKC-Dependent Recruitment of MMP-2 Controls Semaphorin-3A Growth-Promoting Effect in Cortical Dendrites

Gu Z, Cui J, Brown S, Fridman R, Mobashery S, Strongin AY, Lipton SA. A highly specific inhibitor of matrix metalloproteinase-9 rescues laminin from proteolysis and neurons from apoptosis in transient focal cerebral ischemia. Journal of Neuroscience. 2005 Jul 6;25(27):6401–8.

Guoiu M, Sheth S, Nemoto M, Walker M, Pouratian N, Ba A, Toga AW. Cortical spreading depression produces long-term disruption of activity-related changes in cerebral blood volume and neurovascular coupling. Journal of biomedical optics. 2005 Jan;10(1):011004.

Gursoy-Ozdemir Y, Qiu J, Matsuoka N, Bolay H, Bermpohl D, Jin H, Wang X, Rosenberg GA, Lo EH, Moskowitz MA. Cortical spreading depression activates and upregulates MMP-9. The Journal of clinical investigation. 2004 May 15;113(10):1447–55.

Hagemann G, Redecker C, Neumann-Haefelin T, Freund HJ, Witte OW. Increased long-term potentiation in the surround of experimentally induced focal cortical infarction. Annals of Neurology: Official Journal of the American Neurological Association and the Child Neurology Society. 1998 Aug;44(2):255–8.

Hartings JA, Shuttleworth CW, Kirov SA, Ayata C, Hinzman JM, Foreman B, Andrew RD, Boutelle MG, Brennan KC, Carlson AP, Dahlem MA. The continuum of spreading depolarizations in acute cortical lesion development: examining Leao’s legacy. Journal of Cerebral Blood Flow & Metabolism. 2017 May;37(5):1571–94.

Heiss WD. The ischemic penumbra: correlates in imaging and implications for treatment of ischemic stroke. Cerebrovascular Diseases. 2011;32(4):307–20.

Hermann DM, Mies G, Hossmann KA. Biochemical changes and gene expression following traumatic brain injury: role of spreading depression. Restorative neurology and neuroscience. 1999 Jan 1;14(2-3):103–8.

Horiguchi T, Snipes JA, Kis B, Shimizu K, Busija DW. Cyclooxygenase-2 mediates the development of cortical spreading depression-induced tolerance to transient focal cerebral ischemia in rats. Neuroscience. 2006 Jan 1;140(2):723–30.

Hossmann KA. Periinfarct depolarizations. Cerebrovascular and brain metabolism reviews. 1996;8(3):195–208.

Huntley GW. Synaptic circuit remodelling by matrix metalloproteinases in health and disease. Nature Reviews Neuroscience. 2012 Nov;13(11):743–57.

Imamura K, Takeshima T, Fusayasu E, Nakashima K. MMIncreased plasma matrix metalloproteinase-9 levels in migraineurs. Headache: The Journal of Head and Face Pain. 2008 Jan;48(1):135–9.

Itatsu K, Sasaki M, Yamaguchi J, Ohira S, Ishikawa A, Ikeda H, Sato Y, Harada K, Zen Y, Sato H, Ohta T. Cyclooxygenase-2 is involved in the up-regulation of matrix metalloproteinase-9 in cholangiocarcinoma induced by tumor necrosis factor-α. The American journal of pathology. 2009 Mar 1;174(3):829–41.

Jabłonka JA. Post stroke plasticity impairment and its depend-ence on interhemispheric interactions. In MEA Meeting 2012 (p. 136).

Jabłonka JA, Burnat K, Witte OW, Kossut M. Remapping of the somatosensory cortex after a photothrombotic stroke: dynamics of the compensatory reorganization. Neuroscience. 2009 Dec.;165(1):90–100.

Jabłonka JA, Kossut M, Witte OW, Liguz-Lecznar M. Experience-dependent brain plasticity after stroke: effect of ibuprofen and poststroke delay. European Journal of Neuroscience. 2012 Sep;36(5):2632–9.

Jabłonka JA, Urbach P, Kazmierczak M, Borzymowska E, Kublik E. Interhemispheric interac-tions contribution in experience dependent plasticity. Neuroscience Meeting Planner. Washington, DC: Society for Neuroscience. 2014. Program No. 246.12/GG26

Jabłonka JA, Witte OW, Kossut M. Photothrombotic infarct impairs experience-dependent plasticity in neighboring cortex. Neuroreport. 2007 Jan 22;18(2):165–9.

Jander S, Schroeter M, Peters O, Witte OW, Stoll G. Cortical spreading depression induces proinflammatory cytokine gene expression in the rat brain. Journal of Cerebral Blood Flow & Metabolism. 2001 Mar;21(3):218–25.

Kaliszewska A, Bijata M, Kaczmarek L, Kossut M. Experience-dependent plasticity of the barrel cortex in mice observed with 2-DG brain mapping and c-Fos: effects of MMP-9 KO. Cerebral Cortex. 2012 Oct 20;22(9):2160–70.

Katagiri T, Hatano N, Aihara M, Kawano H, Okamoto M, Liu Y, Izumi T, Maekawa T, Nakamura S, Ishihara T, Shirai M. Proteomic analysis of proteins expressing in regions of rat brain by a combination of SDS-PAGE with nano-liquid chromatography-quadrupole-time of flight tandem mass spectrometry. Proteome science. 2010 Dec;8(1):41.

Kaufmann D, Theriot JJ, Zyuzin J, Service CA, Chang JC, Tang YT, Bogdanov VB, Multon S, Schoenen J, Ju YS, Brennan KC. Heterogeneous incidence and propagation of spreading depolarizations. Journal of Cerebral Blood Flow & Metabolism. 2017 May;37(5):1748–62.

Kirkby NS, Chan MV, Zaiss AK, Garcia-Vaz E, Jiao J, Berglund LM, Verdu EF, Ahmetaj-Shala B, Wallace JL, Herschman HR, Gomez MF. Systematic study of constitutive cyclooxygenase-2 expression: role of NF-κB and NFAT transcriptional pathways. Proceedings of the National Academy of Sciences. 2016 Jan 12;113(2):434–9.

Koistinaho J, Chan PH. Spreading depression-induced cyclooxygenase-2 expression in the cortex. Neurochemical research. 2000 May 1;25(5):645–51.

Koistinaho J, Pasonen S, Yrjanheikki J, Chan PH. Spreading depression-induced gene expression is regulated by plasma glucose. Stroke. 1999 Jan 1;30(1):114–9.

Kriz J, Lalancette-Hébert M. Inflammation, plasticity and real-time imaging after cerebral ischemia. Acta neuropathologica. 2009 May 1;117(5):497–509.

Küry P, Schroeter M, Jander S. Transcriptional response to circumscribed cortical brain ischemia: spatiotemporal patterns in ischemic vs. remote non-ischemic cortex. European Journal of Neuroscience. 2004 Apr;19(7):1708–20.

Lauritzen M, Strong AJ. ‘Spreading depression of Leão’and its emerging relevance to acute brain injury in humans. Journal of Cerebral Blood Flow & Metabolism. 2017 May;37(5):1553–70.

Li P, Rudolph U, Huntsman MM. Long-term sensory deprivation selectively rearranges functional inhibitory circuits in mouse barrel cortex. Proceedings of the National Academy of Sciences. 2009 Jul 21;106(29):12156–61.

Li S, Overman JJ, Katsman D, Kozlov SV, Donnelly CJ, Twiss JL, Giger RJ, Coppola G, Geschwind DH, Carmichael ST. An age-related sprouting transcriptome provides molecular control of axonal sprouting after stroke. Nature neuroscience. 2010 Dec;13(12):1496.

Marik SA, Yamahachi H, McManus JN, Szabo G, Gilbert CD. Axonal dynamics of excitatory and inhibitory neurons in somatosensory cortex. PLoS biology. 2010 Jun 15;8(6):e1000395.

Meighan SE, Meighan PC, Choudhury P, Davis CJ, Olson ML, Zornes PA, Wright JW, Harding JW. Effects of extracellular matrix-degrading proteases matrix metalloproteinases 3 and 9 on spatial learning and synaptic plasticity. Journal of neurochemistry. 2006 Mar;96(5):1227–41.

Michaluk P, Wawrzyniak M, Alot P, Szczot M, Wyrembek P, Mercik K, Medvedev N, Wilczek E, De Roo M, Zuschratter W, Muller D. Influence of matrix metalloproteinase MMP-9 on dendritic spine morphology. J Cell Sci. 2011 Oct 1;124(19):3369–80.

Michelson NJ, Kozai TD. Isoflurane and ketamine differentially influence spontaneous and evoked laminar electrophysiology in mouse V1. Journal of neurophysiology. 2018 Aug 1;120(5):2232–45.

Miettinen S, Fusco FR, Yrjänheikki J, Keinänen R, Hirvonen T, Roivainen R, Närhi M, Hökfelt T, Koistinaho J. Spreading depression and focal brain ischemia induce cyclooxygenase-2 in cortical neurons through N-methyl-D-aspartic acid-receptors and phospholipase A2. Proceedings of the National Academy of Sciences. 1997 Jun 10;94(12):6500–5.

Mohajerani MH, Aminoltejari K, Murphy TH. Targeted mini-strokes produce changes in interhemispheric sensory signal processing that are indicative of disinhibition within minutes. Proceedings of the National Academy of Sciences. 2011 May 31;108(22):E183–91.

Mun-Bryce, S., Roberts, L. J. M., Hunt, W. C., Bartolo, A., & Okada, Y. (2004). Acute changes in cortical excitability in the cortex contralateral to focal intracerebral hemorrhage in the swine. Brain Research, 1026(2), 218–226.

Nagel S, Sandy JD, Meyding-Lamade U, Schwark C, Bartsch JW, Wagner S. Focal cerebral ischemia induces changes in both MMP-13 and aggrecan around individual neurons. Brain research. 2005 Sep 14;1056(1):43–50.

Nedergaard M, Hansen AJ. Spreading depression is not associated with neuronal injury in the normal brain. Brain research. 1988 May 24;449(1-2):395–8.

Neumann-Haefelin T, Witte OW. Periinfarct and remote excitability changes after transient middle cerebral artery occlusion. Journal of Cerebral Blood Flow & Metabolism. 2000 Jan;20(1):45–52.

Nonose Y, Gewehr PE, Almeida RF, da Silva JS, Bellaver B, Martins LA, Zimmer ER, Greggio S, Venturin GT, Da Costa JC, Quincozes-Santos A. Cortical bilateral adaptations in rats submitted to focal cerebral ischemia: emphasis on glial metabolism. Molecular neurobiology. 2018 Mar 1;55(3):2025–41.

Overman JJ, Clarkson AN, Wanner IB, Overman WT, Eckstein I, Maguire JL, Dinov ID, Toga AW, Carmichael ST. A role for ephrin-A5 in axonal sprouting, recovery, and activity-dependent plasticity after stroke. Proceedings of the National Academy of Sciences. 2012 Aug 14;109(33):E2230–9.

Que M, Schiene K, Witte OW, Zilles K. Widespread up-regulation of N-methyl-D-aspartate receptors after focal photothrombotic lesion in rat brain. Neurosci Lett. 1999 Oct 1;273(2):77–80.

Piilgaard H, Lauritzen M. Persistent increase in oxygen consumption and impaired neurovascular coupling after spreading depression in rat neocortex. Journal of Cerebral Blood Flow & Metabolism. 2009 Sep;29(9):1517–27.

Potworowski J, Jakuczun W, Łęski S, Wójcik DK. Kernel current source density method. BMC neuroscience. 2011 Dec;12(1):P375.

Redecker C, Wang W, Fritschy JM, Witte OW. Widespread and long-lasting alterations in GABAA-receptor subtypes after focal cortical infarcts in rats: mediation by NMDA-dependent processes. Journal of Cerebral Blood Flow & Metabolism. 2002 Dec;22(12):1463–75.

Reinhard SM, Razak K, Ethell IM. A delicate balance: role of MMP-9 in brain development and pathophysiology of neurodevelopmental disorders. Frontiers in cellular neuroscience. 2015 Jul 29;9:280.

Romanic AM, White RF, Arleth AJ, Ohlstein EH, Barone FC. Matrix metalloproteinase expression increases after cerebral focal ischemia in rats. Stroke. 1998 May 1;29(1020):30.

Samad TA, Moore KA, Sapirstein A, Billet S, Allchorne A, Poole S, Bonventre JV, Woolf CJ. Interleukin-1β-mediated induction of Cox-2 in the CNS contributes to inflammatory pain hypersensitivity. Nature. 2001 Mar;410(6827):471.

Sang N, Chen C. Lipid signaling and synaptic plasticity. The Neuroscientist. 2006 Oct;12(5):425–34.

Schiene K, Bruehl C, Zilles K, Qu M, Hagemann G, Kraemer M, Witte OW. Neuronal hyperexcitability and reduction of GABAA-receptor expression in the surround of cerebral photothrombosis. Journal of Cerebral Blood Flow & Metabolism. 1996 Sep;16(5):906–14.

Shinohara M, Dollinger B, Brown G, Rapoport S, Sokoloff L. Cerebral glucose utilization: local changes during and after recovery from spreading cortical depression. Science. 1979 Jan 12;203(4376):188–90.

Sommer CJ. Ischemic stroke: experimental models and reality. Acta neuropathologica. 2017 Feb 1;133(2):245–61.

Song M, Yu SP. Ionic regulation of cell volume changes and cell death after ischemic stroke. Transl Stroke Res. 2014 Feb;5(1):17–27.

Strominger RN, Woolsey TA. Templates for locating the whisker area in fresh flattened mouse and rat cortex. Journal of neuroscience methods. 1987 Dec 1;22(2):113–8.

de Souza TK, e Silva MB, Gomes AR, de Oliveira HM, Moraes RB, de Freitas Barbosa CT, Guedes RC. Potentiation of spontaneous and evoked cortical electrical activity after spreading depression: in vivo analysis in well-nourished and malnourished rats. Experimental brain research. 2011 Oct 1;214(3):463–9.

Tailby C, Wright LL, Metha AB, Calford MB. Activity-dependent maintenance and growth of dendrites in adult cortex. Proceedings of the National Academy of Sciences. 2005 Mar 22;102(12):4631–6.

Theriot JJ, Toga AW, Prakash N, Ju YS, Brennan KC. Cortical sensory plasticity in a model of migraine with aura. Journal of Neuroscience. 2012 Oct 31;32(44):15252–61.

Urbach A, Bruehl C, Witte OW. Microarray-based long-term detection of genes differentially expressed after cortical spreading depression. European Journal of Neuroscience. 2006 Aug;24(3):841–56.

Urbach A, Brueckner J, Witte OW. Cortical spreading depolarization stimulates gliogenesis in the rat entorhinal cortex. Journal of Cerebral Blood Flow and Metabolism. 2014 Dec 35(4), 576–582.

Vecchia D, & Pietrobon D. Migraine: a disorder of brain excitatory – inhibitory balance? Trends in Neurosciences. 2012 Aug; 35(8), 507–520.

Victor NA, Wanderi EW, Gamboa J, Zhao X, Aronowski J, Deininger K, Lust WD, Landreth GE, Sundararajan S. Altered PPARγ expression and activation after transient focal ischemia in rats. European Journal of Neuroscience. 2006 Sep 24(6):1653–63.

Wang X, Jung J, Asahi M, Chwang W, Russo L, Moskowitz MA, Dixon CE, Fini ME, Lo EH. Effects of matrix metalloproteinase-9 gene knock-out on morphological and motor outcomes after traumatic brain injury. Journal of Neuroscience. 2000 Sep 15;20(18):7037–42.

Wang XB, Bozdagi O, Nikitczuk JS, Zhai ZW, Zhou Q, Huntley GW. Extracellular proteolysis by matrix metalloproteinase-9 drives dendritic spine enlargement and long-term potentiation coordinately. Proceedings of the National Academy of Sciences. 2008 Dec 9;105(49):19520–5.

Witte OW, Bidmon HJ, Schiene K, Redecker C, Hagemann G. Functional differentiation of multiple perilesional zones after focal cerebral ischemia. Journal of Cerebral Blood Flow & Metabolism. 2000 Aug;20(8):1149–65.

Woessner Jr JF. Matrix metalloproteinases and their inhibitors in connective tissue remodeling. The FASEB Journal. 1991 May;5(8):2145–54.

Yin W, Badr AE, Mychaskiw G, Zhang JH. Down regulation of COX-2 is involved in hyperbaric oxygen treatment in a rat transient focal cerebral ischemia model. Brain Res. 2002 Feb 1;926(1-2):165–71.

Yin KJ, Deng Z, Hamblin M, Zhang J, Chen YE. Vascular PPARδ protects against stroke-induced brain injury. Arteriosclerosis, thrombosis, and vascular biology. 2011 Mar;31(3):574–81.

